# EphB1 in Endothelial Cells Regulates Caveolae Formation and Endocytosis

**DOI:** 10.1101/619510

**Authors:** Chinnaswamy Tiruppathi, Sushil C. Regmi, Dong-Mei Wang, Gary C.H. Mo, Peter T. Toth, Stephen M. Vogel, Radu V. Stan, Mark Henkemeyer, Richard D. Minshall, Asrar B. Malik

## Abstract

Caveolae, the cave-like structures abundant in endothelial cells (ECs), are important in regulating key functions such as caveolae-mediated endocytosis and generation of nitric oxide. Here we show that deletion of the receptor tyrosine kinase EphB1 (*EphB1*^−/−^) in mice markedly reduced the caveolae number in ECs of heart and lung vessels and prevented caveolae-mediated endocytosis. EphB1 expressed in adult ECs was shown to bind the caveolin-1 (Cav-1) scaffold domain (CSD) via the CSD binding motif (CSDBM) on EphB1. We demonstrated that activation of EphB1 by the native ligand Ephrin B1 uncoupled EphB1 from Cav-1, and licensed *Src-*dependent Y-14 Cav-1 phosphorylation. Deletion of CSDBM on EphB1 prevented EphB1/Cav-1 interaction and the activation of *Src* and *Src* mediated Y-14 Cav-1 phosphorylation. These studies identify the central role of endothelium expressed EphB1 in regulating caveolae biogenesis and caveolae-mediated endocytosis.

## Introduction

The <u>**e**</u>rythropoietin-<u>**p**</u>roducing <u>**h**</u>epatocellular carcinoma (Eph) receptor tyrosine kinases and their ligands (Ephrins) are expressed multiple cell types (1,2). Ephrins have important roles in regulating neuronal and vascular development, axon guidance, and maintenance of cell-specific identities (3–6). While the functions of Ephrins are known in the embryo, their molecular understanding in adult cells is less clear (1–6). Here based on evidence of the key role of Ephrins in vascular development, postnatal angiogenesis, and vascular integrity (5–9), we investigated whether they can be redeployed in adult endothelial cells (EC) to regulate vascular function.

Caveolae are plasma membrane invaginations in ECs (10–12) in which caveolin-1 (Cav-1) is a primary structural and signaling protein inserted in the caveolar membrane (11–14). Both Cav-1 and the isoform Cav-2 are expressed in ECs (12, 15) and low-affinity electrostatically controlled protein-protein interactions between the two enables the formation of caveolae (15–18). Caveolae on the luminal side of ECs have several functions (16) among them loading of cargo and caveolae-mediated endocytosis (17). Deletion of Cav-1 (*Cav-1*^−/−^) in mice showed flattening of caveolae indicating the importance of Cav-1 for the formation of caveolae in ECs (19,20). Further, these mice failed internalize tracer proteins in contrast to controls (19,20). We showed that *Src* kinase-mediated phosphorylation of Cav-1 on Y-14 was required for fission of caveolae from the plasma membrane and caveolae-mediated endocytosis (21–23). Cav-1 scaffold domain (CSD) binds other proteins via a peptide sequence in CSD-interacting proteins, the CSD binding motif (CSDBM) (Fig. 1A). Several of these key binding proteins have been identified including *Src* tyrosine kinases, eNOS (endothelial nitric oxide synthase), and trimeric G protein subunits, Ras, and PPARγ (24–33). Interestingly, in a study using CHO cells, it was shown that EphB1 can also bind CSD via its CSDBM (^808^WSYGIVMW^815^) and target the plasma membrane (34); however, while this study raised the potential for this interaction it did not address the mechanism of this interaction and in functional significance in the endothelium.

**Figure 1.**
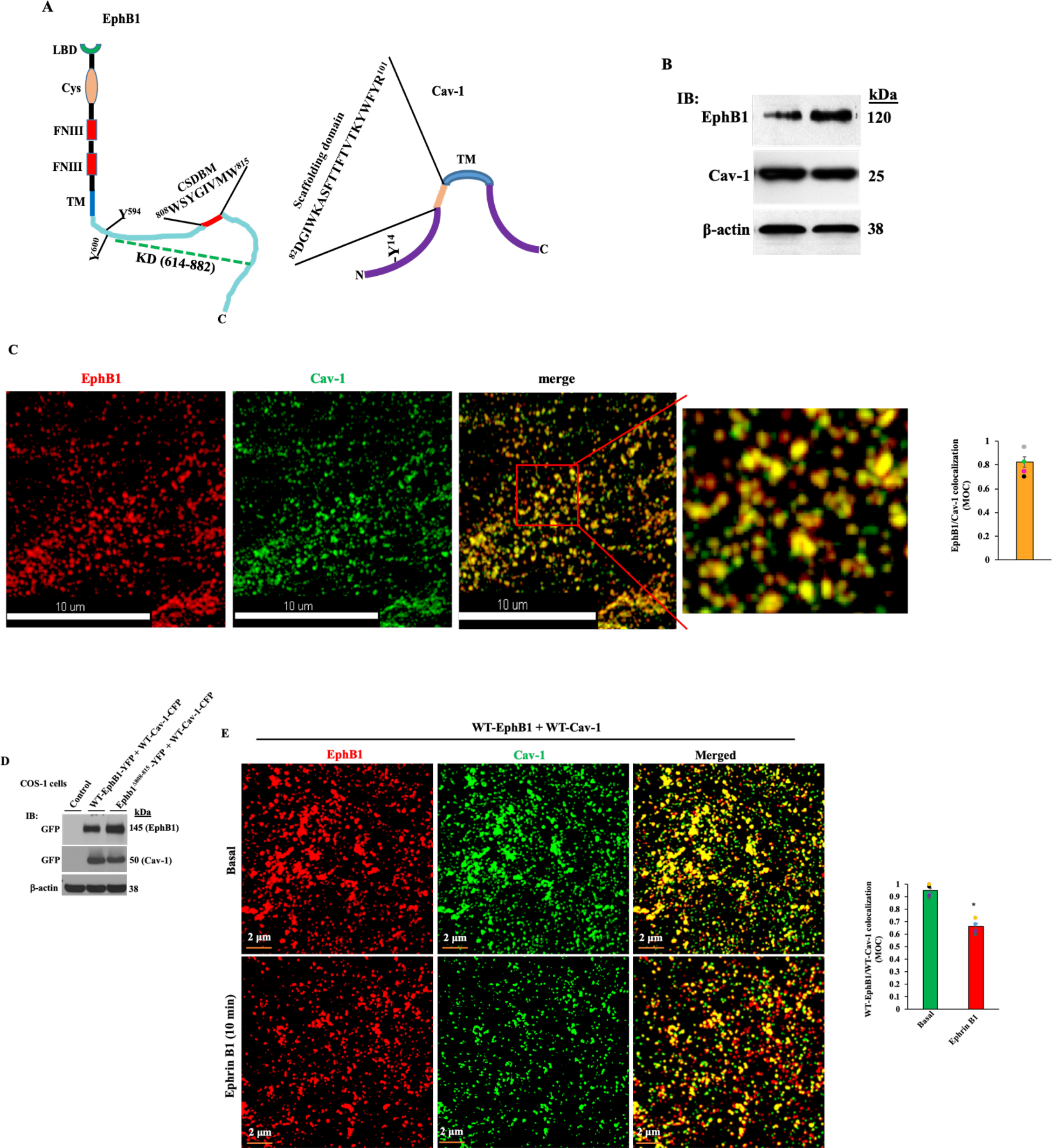
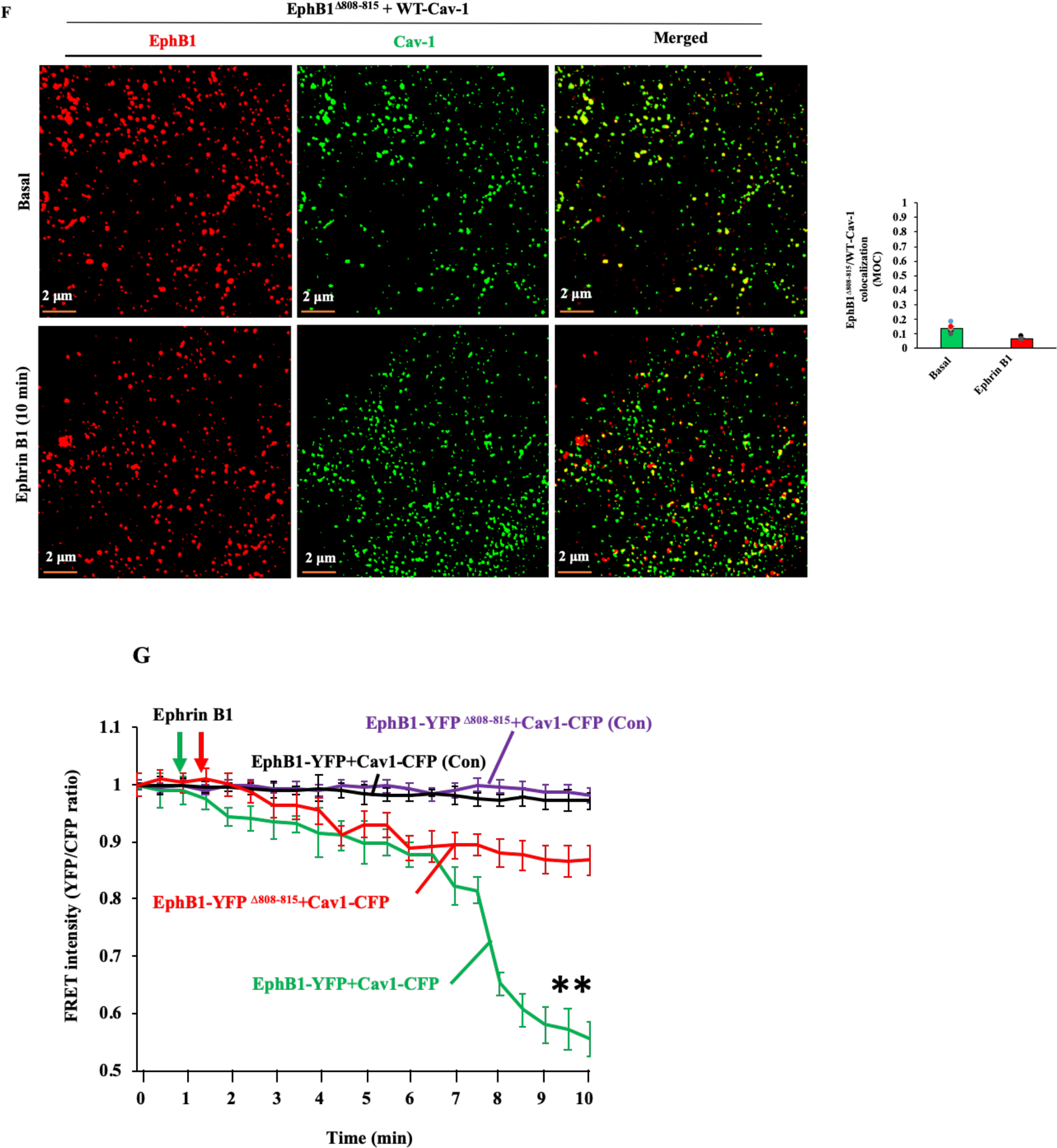
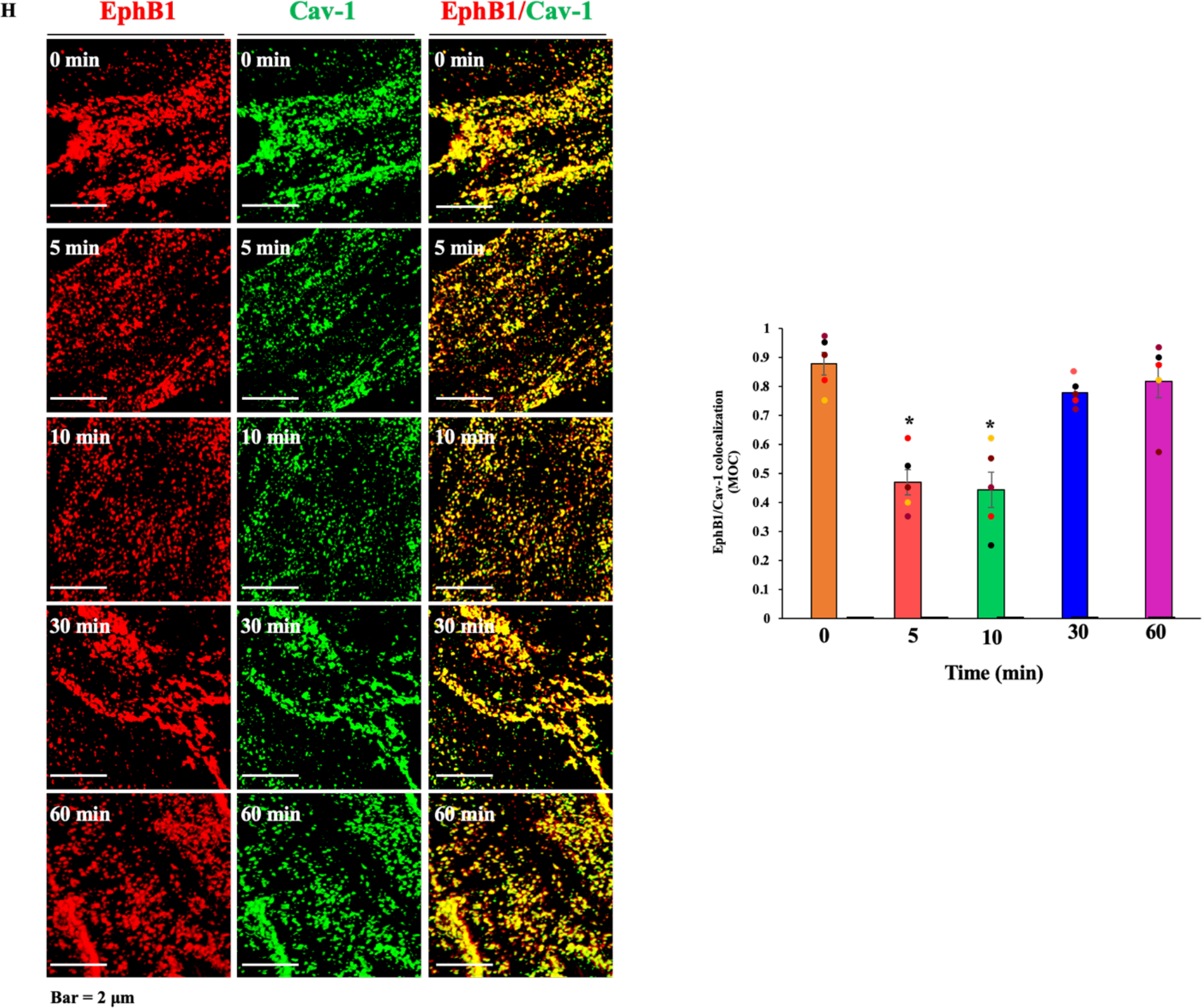
EphB1 interacts with the caveolin-1 (Cav-1) scaffold binding domain (CSD). **(A)** Schematic representation of structural domain features of EphB1 and Cav-1. CSD, Cav-1 scaffold domain (CSD); CSDBM, CSD binding motif in EphB1. LBD, ligand-binding domain; Cys, cysteine-rich domain; FNIII, fibronectin-type III repeats; TM, transmembrane domain; KD (dashed green line), EphB1 kinase domain (614–882). **(B)** Western analysis of human lung microvessel endothelial cells (HLMVECs) showing expression of EphB1 and Cav-1. **(C)** Sectional view of single cell plasma membrane image from 3D-structured illumination microscopy (SIM) showing co-localization of EphB1 with Cav-1 in HLMVECs. *In merge*, magnified view of the region indicated shown in right. Co-localization efficiency between EphB1 and Cav-1 as assessed by MOC is shown in *right panel*. N = 4 cells. **(D)** Western analysis of COS-1 cells transfected with WT-EphB1-YFP (WT-EphB1 C-terminus fused with YFP), WT-Cav-1-CFP (WT-Cav-1 C-terminus fused with CFP), and EphB1^Δ808-815^-YFP. **(E)** Sectional image of COS-1 cell expressing WT-EphB1 plus WT-Cav-1 using 3D-SIM showing CSDBM of EphB1 interacts with WT-Cav-1. *Top panel*, representative unstimulated cell sectional view of 3D image. *Bottom panel*, representative COS-1 cell sectional view of 3D image showing effect of Ephrin B1-Fc (1 μg/ ml) stimulation which caused EphB1 dissociation from Cav-1. *Right panel*, EphB1 and Cav-1 co-localization efficiency assessed by MOC. N = 4 cells/group; *p< 0.5, compared with basal. **(F)** Sectional image of COS-1 cell expressing EphB1^Δ808-815^ plus WT-Cav-1 using 3D-SIM showing absence of interaction between CSDBM deleted EphB1 (EphB1^Δ808-815^) and WT-Cav-1. *Top panel*, control. *Bottom Panel*, stimulated with Ephrin B1-Fc (1 μg/ml). *Right panel*, colocalization efficiency between EphB1^Δ808-815^ and Cav-1 assessed by MOC. **(G)** Live cell FRET measurements in COS-1 cells showing dissociation of EphB1 from Cav-1 after challenging with Ephrin B1. YFP/CFP ratio normalized to 1 before challenging cells with the ligand Ephrin B1 (Ephrin B1-Fc;1 μg/ ml). Results shown are mean ± SE of 3 separate experiments. **p< 0.001, compared with WT-EphB1 + WT-Cav-1 control (green) or EphB1^Δ808-815^ WT-Cav-1 (red) exposed to Ephrin B1. **(H)** Sectional images of single WT-endothelial cells (ECs) using 3D-SIM showing changes in co-localization of EphB1 with Cav-1 at baseline and following stimulation with the ligand Ephrin B1-Fc. ECs from WT mice were serum starved for 2h and then exposed to Ephrin B1-Fc (1μg/ml) for different times up to 60 min, and immune-stained with antibodies specific to EphB1 and Cav-1. *Right panel* shows the EphB1 and Cav-1 co-localization efficiency assessed by MOC. N = 5 cells/group; *p< 0.5, compared with basal.

In the present study, using mouse models and endothelial cells, we showed a crucial function of EphB1 in regulating the genesis of caveolae in ECs. Deletion of EphB1 in mice severely depleted caveolae numbers. Activation of EphB1 with the ligand Ephrin B1 disassociated EphB1 from Cav-1 to induce EphB1 auto-phosphorylation on Y-600 via EphB1’s kinase domain and thereby recruited *Src*. We demonstrated that *Src* activation secondary to its recruitment induced Cav-1 phosphorylation on Y-14 and was required for the initiation of caveolae-mediated endocytosis. Thus, our results demonstrate an intersection between EphB1 and Cav-1 through a binding interaction that regulates the biogenesis and function of caveolae in ECs important in vascular homeostasis.

## Results

### EphB1 binds the Cav-1 scaffold domain in endothelial cells

Studies were made in human lung microvascular ECs (HLMVECs) prominently expressing both EphB1 (Fig. 1A) and Cav-1 (Fig. 1B). Using 3D-SIM super-resolution microscopy, we observed an interaction between EphB1 and Cav-1 as measured by Manders overlap coefficient (MOC) (35) (Fig. 1C). In unstimulated cells, both EphB1 and Cav-1 were clustered in multilobed caveolar rosettes (Fig. 1C). In other studies, co-expression of EphB1 and Cav-1 in COS-1 cells also showed a strong interaction (Fig. 1D-E; **Supplemental Figure 1A and B**). Using the mutant EphB1^Δ808-815^ in which CSDBM is deleted, we observed that it did not bind WT-Cav-1 (Fig. 1F; **Supplemental Figure 1A and B**).

To determine whether activation of EphB1 influences EphB1/Cav-1 interaction, we added Ephrin B1-Fc to COS-1 cells co-expressing EphB1 and Cav-1. Soluble Ephrin extracellular domain fused to Fc domain (Ephrin-Fc), a standard Eph receptor agonist (1,2), was added. Ephrin B1-Fc significantly reduced EphB1/Cav-1 interaction (Fig. 1E; **Supplemental Figure 1A and B)**. To further study this interaction, we also co-expressed WT-EphB1-YFP with either WT-Cav-1-CFP or mutant EphB1^Δ808-815^-YFP with WT-Cav-1-CFP in COS-1 cells, and measured live cell FRET efficiency (23). Cells co-expressing WT-EphB1-YFP and WT-Cav-1-CFP activated by Ephrin B1-Fc showed markedly reduced FRET intensity (Fig. 1G) whereas cells co-expressing EphB1^Δ808-815^-YPF and WT-Cav-1-CFP showed no significant response (Fig. 1G). We also determined EphB1/Cav-1 interaction in murine lung ECs (C57BL/6 mice) by immunostaining using antibodies to EphB1 and Cav-1, and observed marked co-localization in caveolae which was markedly reduced within minutes of adding Ephrin B1 with the effect reversed over time (Fig. 1 H; **Supplemental Figure 1C and D)** as described in **Supplemental Figure 1E**.

### EphB1/Cav-1 interaction occludes *Src* activation and Y-14 Cav-1 phosphorylation

EphB1 contains 2 tyrosine residues (Y^594^ and Y^600^) in its juxta-membrane region (1,2,7,36). Ephrin B1 activation of EphB1 is known to induce auto-phosphorylation of EphB1 on Y^600^, which thereby binds *Src* homology-2 (SH2) domain of *Src* family kinases (SFKs) to activate SFKs (36). To address whether EphB1 contributes *Src-*dependent phosphorylation of Cav-1 on Y-14, we measured Ephrin B1-induced EphB1 autophosphorylation, *Src* activation, and Y-14 phosphorylation of Cav-1 in HLMVECs. We observed that Ephrin B1 induced EphB1 auto-phosphorylation on Y^600^, phosphorylation of *Src* on Y-416 as well as Cav-1 on Y-14 (Fig. 2, B-D). Since other EphB receptors may also be activated by Ephrin B1, we addressed whether the effects of the ligand in ECs were specific to EphB1. Thus, we used an EphB1 selective antagonistic peptide (EphB1-A-pep) (Fig. 2A) (37,38), which prevents binding of Ephrin B1 to EphB1 ligand binding domain and blocks EphB1 signaling (37,38). HLMVECs treated with EphB1-A-pep prevented Ephrin B1-induced EphB1 auto-phosphorylation (Fig. 2B), *Src* activation (p-Y416) (Fig. 2C), and pY-14 Cav-1 phosphorylation as compared to a control peptide (Fig. 2D).

**Figure 2.**
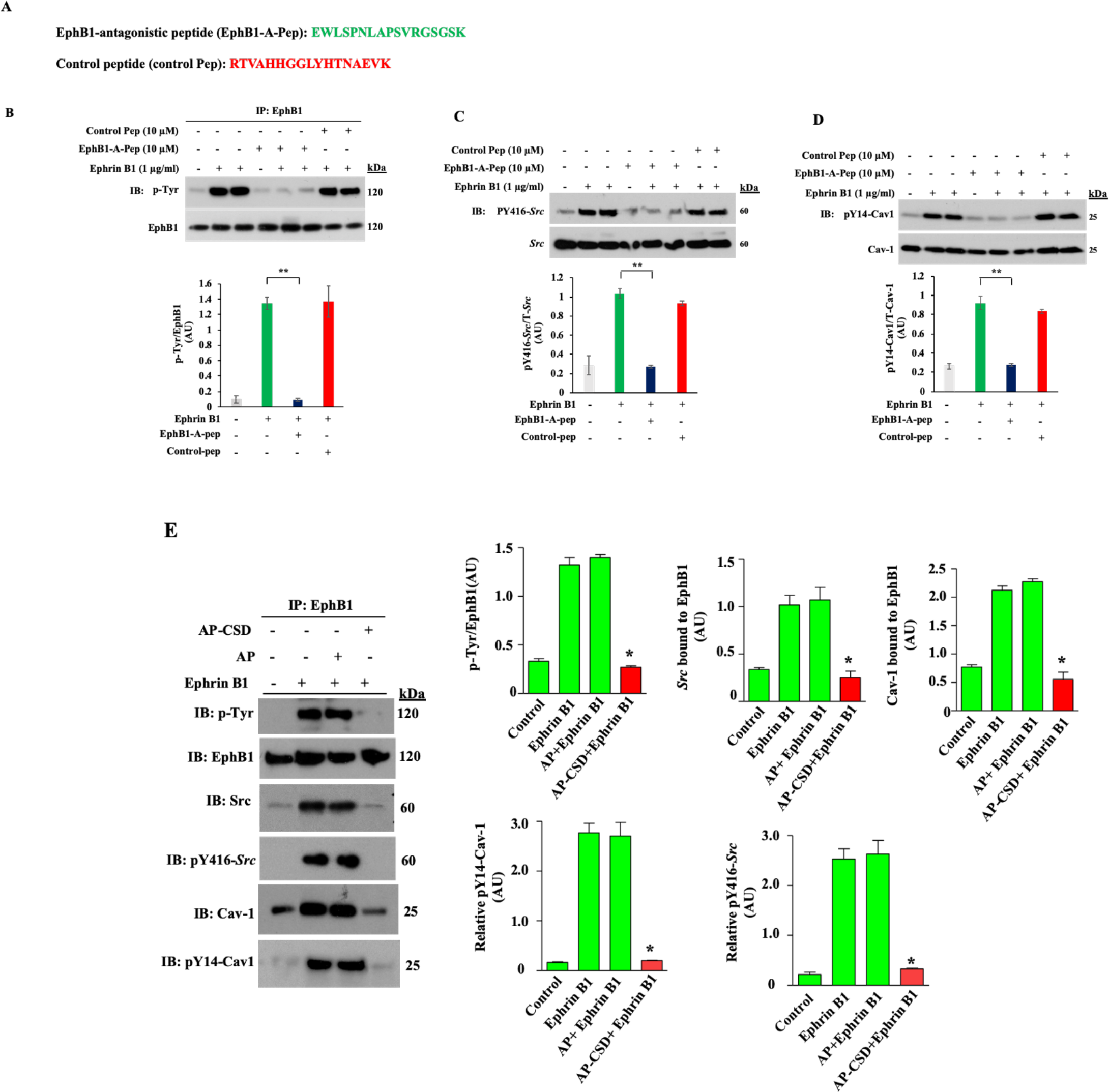
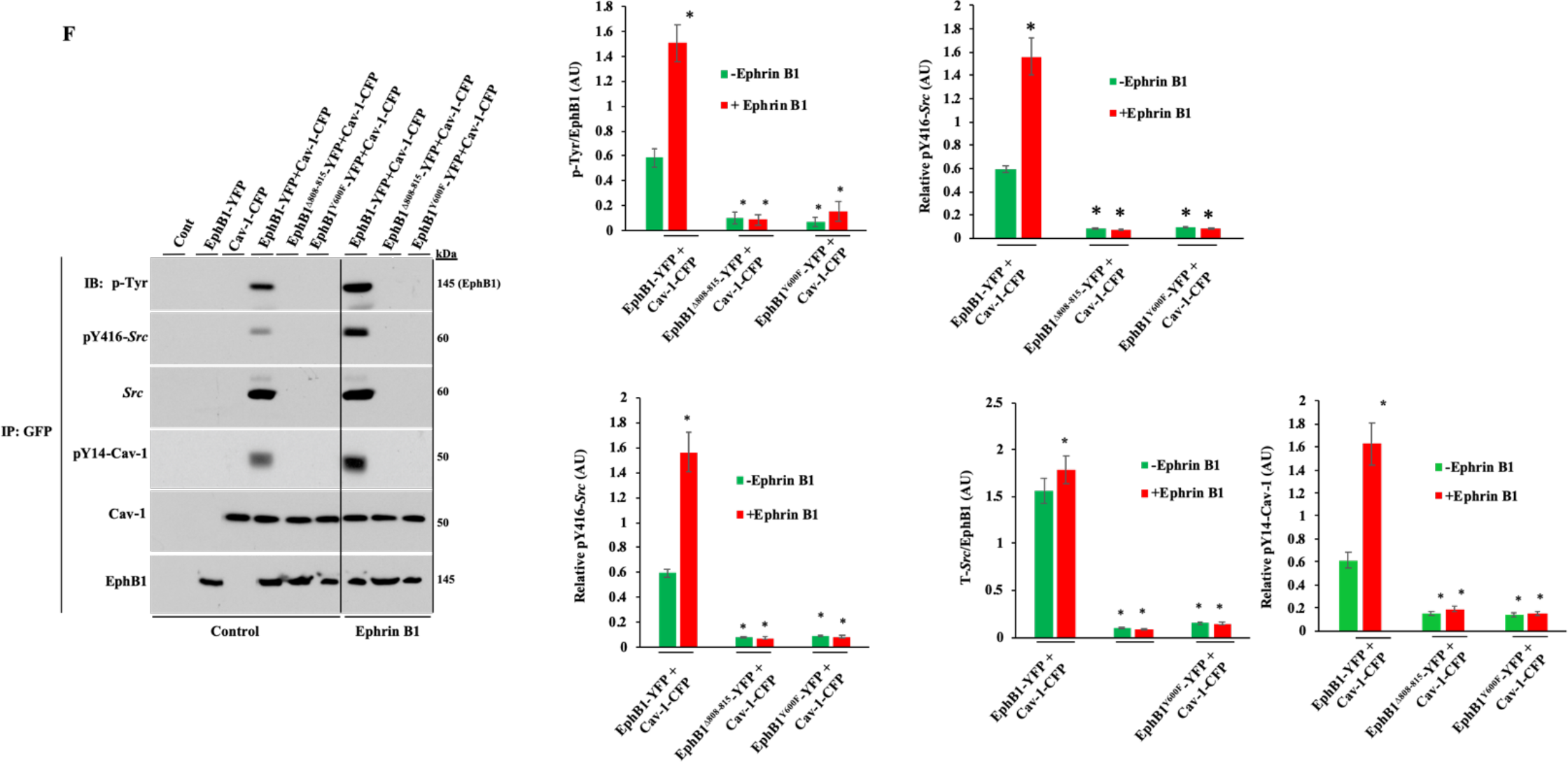
(A-D) EphB1 specific antagonistic peptide prevents Ephrin B1-induced auto-phosphorylation of EphB1, *Src* activation, and phosphorylation of Cav-1 on Y-14. **(A)** Sequence of EphB1 antagonistic peptide (EphB1-A-Pep) and control peptide (Control Pep). (**B-D)** HLMVECs incubated in serum free condition for 2 h at 37°C were treated with EphB1-Ap-pep or control peptide (Control-pep) for 30 min. Cells were then exposed to EphrinB1 (EphrinB1-Fc) (1µg/ml) for 10 min at 37°C. **(B)** Cell lysates immunoprecipitated with anti-EphB1 pAb and blotted with anti-phospho-tyrosine mAb to determine phosphorylation of EphB1. **(C)** Total cell lysates were used to determine phosphorylation of *Src* at Y416 to assess *Src* activation. **(D)** Total cell lysates were used to determine phosphorylation of Cav-1 on Y-14. (**B-D)**, blots shown are representative of 3 separate experiments. **p<0.001, Control *vs* EphB1-A-pep. **(E) Cell permeable CSD peptide prevents Ephrin B1-induced autophosphorylation of EphB1, *Src* activation, and Cav-1 phosphorylation on Y-14.** HLMVECs incubated with serum free medium for 2 h at 37°C were incubated with cell permeable control AP peptide (5µM) or AP-CSD peptide (5 µM) for 60min at 37°C. Cells were then stimulated with Ephrin B1-Fc (1µg/ml) for 10min at 37°C. Cell lysates IP-ed with anti-EphB1 pAb and IP-ed samples were immunoblotted with the indicated antibodies. Results shown are representative of 3 experiments. *p<0.05, compared with controls. **(F) Binding of CSDBM of EphB1 to CSD and phosphorylation EphB1 on Y-600 are required *Src* activation and Y-14 Cav-1 phosphorylation.** WT-EphB1 + WT-Cav-1, EphB1^Δ808-815^ WT-Cav-1, or EphB1^Y600F^ + WT-Cav-1 expressing Cos-1 cells were stimulated with or without EphrinB1-Fc (1µg/ml) for 10 min. Cell lysates were immunoprecipitated with anti-GFP mAb (anti-GFP mAb immunoprecipitates both YFP-tagged EphB1 and CFP-tagged Cav-1) were blotted with indicated antibodies. Results shown are representative of 3 experiments. **p < 0.001.

We next determined whether the cell-permeable synthetic CSD (antennapedia (AP)-CSD) peptide can also prevent EphB1/Cav-1 interaction and downstream *Src* activation. Ephrin B1-Fc stimulation caused auto-tyrosine phosphorylation of EphB1, *Src* binding to EphB1, and phosphorylation of *Src* at Y-416 and of Cav-1 at Y-14 in control AP peptide-treated HLMVECs whereas these responses in AP-CSD peptide-loaded cells were blocked (Fig. 2E).

To define further EphB1-mediated *Src* activation and Cav-1 phosphorylation, we co-expressed WT-EphB1-YFP, mutant EphB1^Δ808-815^-YFP, or mutant EphB1^Y600F^-YFP with WT-Cav-1-CFP in COS-1 cells. In WT-EphB1- and WT-Cav-1 co-expressing cells, we observed auto-phosphorylation of EphB1 and phosphorylation of both *Src* on Y-416 and Cav-1 on Y-14, which were both increased by the Ephrin B1-Fc ligand (Fig. 2F). In EphB1^Δ808-815^ and WT-Cav-1 co-expressing cells, we observed inhibition of basal as well as Ephrin B1-induced auto-phosphorylation of EphB1 as well as phosphorylation of *Src* and Cav-1 (Fig. 2F). Similar results were seen in mutant EphB1^Y600F^ and WT-Cav-1 co-expressing cells (Fig. 2F). Thus, the interaction of CSD and CSDBM is required for Ephrin B1-induced auto-phosphorylation of EphB1 on Y-600, EphB1-mediated *Src* activation, and Cav-1 phosphorylation on Y-14.

### EphB1 regulates the biogenesis of caveolae

We next studied mice in which EphB1 was deleted (*EphB1*^−/−^) and in mice expressing EphB1-tc (EphB1-βgal fusion receptor lacking the tyrosine kinase and C-terminal domains). Western blot analysis of lung tissue and isolated lung ECs showed the absence of EphB1 expression in *EphB1*^−/−^ mice (**Supplemental Figure 2, A and B**) and expression of EphB1-βgal fusion receptor in EphB1-tc mice (**Supplemental Figure 2, A and B**). Immunostaining of lung sections from EphB1-tc mice with antibodies specific to β-gal and vWF (an EC marker) showed β-gal expression in blood vessels (**Supplemental Figure 2C).** As EphB1 interacts with Cav-1 as shown above, using ECs obtained from *EphB1*^−/−^ mice we determined whether EphB1 is involved in regulating Cav-1 expression, and found that Cav-1 protein expression was markedly reduced in *EphB1*^−/−^ mice as compared to WT (Fig. 3A); however, decreased Cav-1 protein expression was not associated with reduced Cav-1 mRNA in *EphB1*^−/−^ mice (Fig. 3B). Since Cav-1 expression may be downregulated by ubiquitination via the proteasomal pathway (39,40), we observed increased ubiquitination of Cav-1 in lung ECs of *EphB1*^−/−^ mice compared to WT (Fig. 3C). We also determined the expression of caveolar coat proteins cavin-1 and -2 important for caveolae biogenesis (41,42). Cavin-1 expression was similar in both WT and *EphB1*^−/−^ mice and Cavin-2 expression was not reduced in *EphB1*^−/−^ ECs (Fig. 3D), ruling out reduced Cavin-1, -2 expression as the basis of defective caveolae biogenesis in *EphB1*^−/−^ mice.

**Figure 3.**
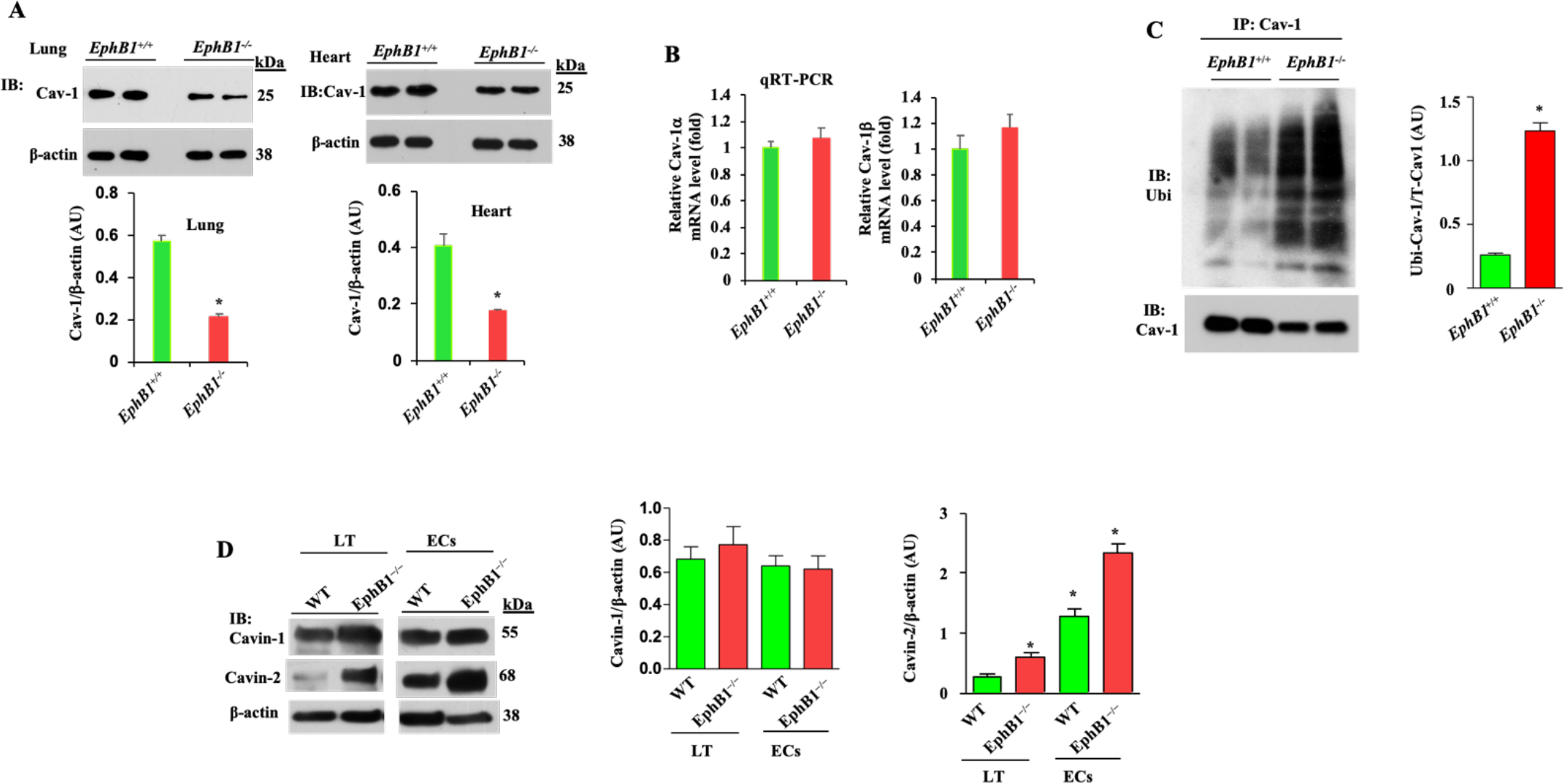
EphB1 regulation of expression of Cav-1. **(A)** Lung and heart tissues from WT (*EphB1*^+/+^) and EphB1 knockout (*EphB1*^−/−^) mice were used for Western blotting to determine Cav-1 expression. n = 5 mice per group; unpaired *t*-test. *p<0.05. **(B)** EphB1 deficiency did not alter Cav-1 mRNA expression. QRT-PCR was performed utilizing total RNA from lung tissue of WT and *EphB1*^−/−^ mice to determine mRNA expression for Cav-1α and Cav-1β isoforms. **(C)** EphB1 deficiency induced Cav-1 degradation via ubiquitination. Freshly isolated lung ECs from WT and *EphB1*^−/−^ mice were IP-ed with anti-Cav-1 mAb and blotted with anti-pan-ubiquitin (Ubi) pAb. n = 5 mice per genotype used for EC isolation; unpaired *t*-test. *p<0.05. **(D)** Expression of caveolar coat proteins Cavin-1 and −2 was not suppressed in *EphB1*^−/−^ mice. Lung tissue and lung ECs (ECs) from WT (*EphB1*^+/+^) and *EphB1*^−/−^ mice were used to determine Cavin-1 and −2 by Western analysis. n = 5 mice per group; 3 separate EC preparations were used. unpaired *t*-test. *p<0.05.

By ultrastructural analysis of caveolae, we confirmed defective caveolar morphogenesis in lung and heart microvascular endothelia of *EphB1*^−/−^ mice (Fig. 4, A and C). The number of caveolae was significantly reduced in ECs of lungs (Fig. 4A, *right panel*) (4.2 ± 0.2/µm in WT mice *vs.* 1.3 ± 0.2/µm *EphB1*^−/−^ mice) and of hearts (5.4 ± 0.3/µm in *EphB1*^+/+^ mice *vs.* 2.6 ± 0.3/µm *EphB1*^−/−^ mice, Fig. 4C, *right panel*). However, there was no difference in the caveolar shape (Fig. 4A and C) or inter-endothelial junctions (IEJs) (Fig. 4B).

**Figure 4.**
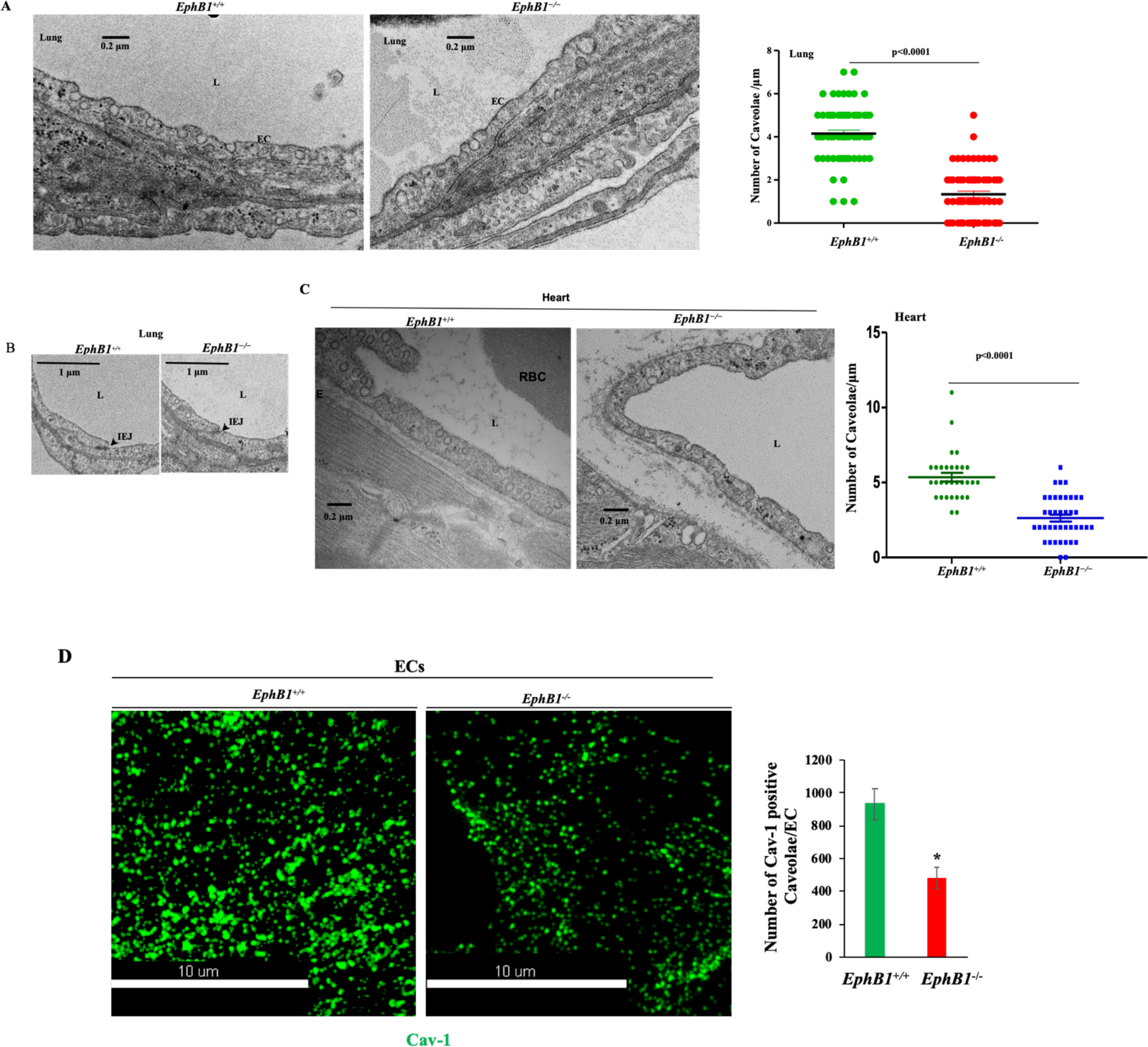
EphB1 regulates caveolae morphogenesis in endothelial cells. (**A**) Electron micrograph showing reduced caveolae number in lung endothelia of *EphB1*^−/−^ mice with morphometric data in (*right panel*). Caveolae open at the cell surface were counted per micrometer of endothelial luminal plasma membrane. Multiple electron micrographs were used for counting caveolae; *EphB1*^+/+^ (WT) lungs, N = 67; *EphB1*^−/−^ lungs, N=66. L, lumen; EC, endothelium; p< 0.0001 vs. *EphB1*^+/+^. (**B**) EphB1 deletion did not alter lung inter-endothelial junctions (IEJ). **(C)** Electron micrograph showing reduced caveolae number in heart endothelia of *EphB1*^−/−^ mice with morphometric data in (*right panel*). *EphB1*^+/+^ (WT) hearts, N = 31; *EphB1*^−/−^ hearts, N=40. L, lumen; p< 0.0001 vs. *EphB1*^+/+^. **(D)**3D-SIM super-resolution microscopy image showing reduced number of Cav-1+ve structures in *EphB1*^−/−^-ECs. ECs from WT and *EphB1*^−/−^ mice were stained with anti-Cav-1 pAb and used to obtain images by 3D-SIM super-resolution microscopy. Representative sectional view of single cell plasma membrane image from 3D-SIM super-resolution microscopy showing Cav-1+ve vesicles in LECs from *EphB1*^+/+^ and *EphB1*^−/−^ mice. N = 5 cells per genotype; *p< 0.05.

Studies were also made using lung ECs from WT and *EphB1*^−/−^ mice in which we determined morphology and function of caveolae (defined as Cav-1+ve structures) using 3D-SIM super-resolution microscopy. Cav-1+ve caveolae number was markedly reduced in ECs of *EphB1*^−/−^ mice and caveolar rosette-like structures in ECs formed by fusion of multiple caveolae were not seen in ECs from *EphB1*^−/−^ mice as compared to WT (Fig. 4D).

### EphB1 is required for caveolae endocytosis

To address the requirement for EphB1 in signaling caveolae mediated endocytosis, we used ECs *EphB1*^−/−^ as well as from *Cav-1*^−/−^, and determined caveolae-mediated endocytosis in response to Ephrin B1. Ephrin B1 induced time-dependent increases in phosphorylation of *Src* on Y-416 and Y-14 Cav-1 in WT ECs whereas these responses were blocked in ECs from *Cav-1*^−/−^ mice (Fig. 5A, B). In ECs from *EphB1*^−/−^ mice, we observed that Ephrin B1-induced phosphorylation on *Src* Y-416 and Cav-1 Y-14 were also blocked (Fig. 5C and D). Using ECs from *EphB1*^−/−^ and *Cav-1*^−/−^ mice, we next addressed whether EphB1/Cav-1 interaction was required for signaling caveolae-mediated endocytosis. Here we determined the endocytic function of caveolae by quantifying the uptake of tracer albumin Alexa Fluor-594 labeled bovine serum albumin tracer (Alexa-594 BSA) (17,23,43) using ECs from *EphB1*^−/−^ and *Cav-1*^−/−^ mice. Studies were made in ECs incubated with serum free medium for 2h followed by addition of Ephrin B1-Fc ligand (1 μg/ml) in medium containing 100 μg/ml albumin tracer plus 2 mg/ml unlabeled albumin. Analysis of single cell 3D images showed a time-dependent albumin internalization in ECs from WT mice as early as 5 min after tracer addition. We observed 16 ± 3 particles/cell with maximum uptake of 850 ± 14 particles/cell attained seen at 60 min (Fig. 6A and Table 1). We also observed that internalized albumin particles trafficked into the cell, ~3 μm from the apical side (Fig. 6A). Numerous Cav-1+ve caveolae were seen in ECs of WT mice (Fig. 6A) whereas ECs from *EphB1*^−/−^ mice showed 4-fold reduction in albumin tracer internalization (in WT-ECs 850 ± 14 particles/cell at 60 min vs. in *EphB1*^−/−^-ECs 216 ± 16 particles/cell) (Fig. 6A, B; Table 1). We similarly measured albumin internalization in ECs of *Cav-1*^−/−^ mice as a comparison with *EphB1*^−/−^-ECs, and also observed inhibition of time-dependent internalization of albumin particles in *Cav-1*^−/−^-ECs (Fig. 6C, D and Table 1).

**Table 1.**
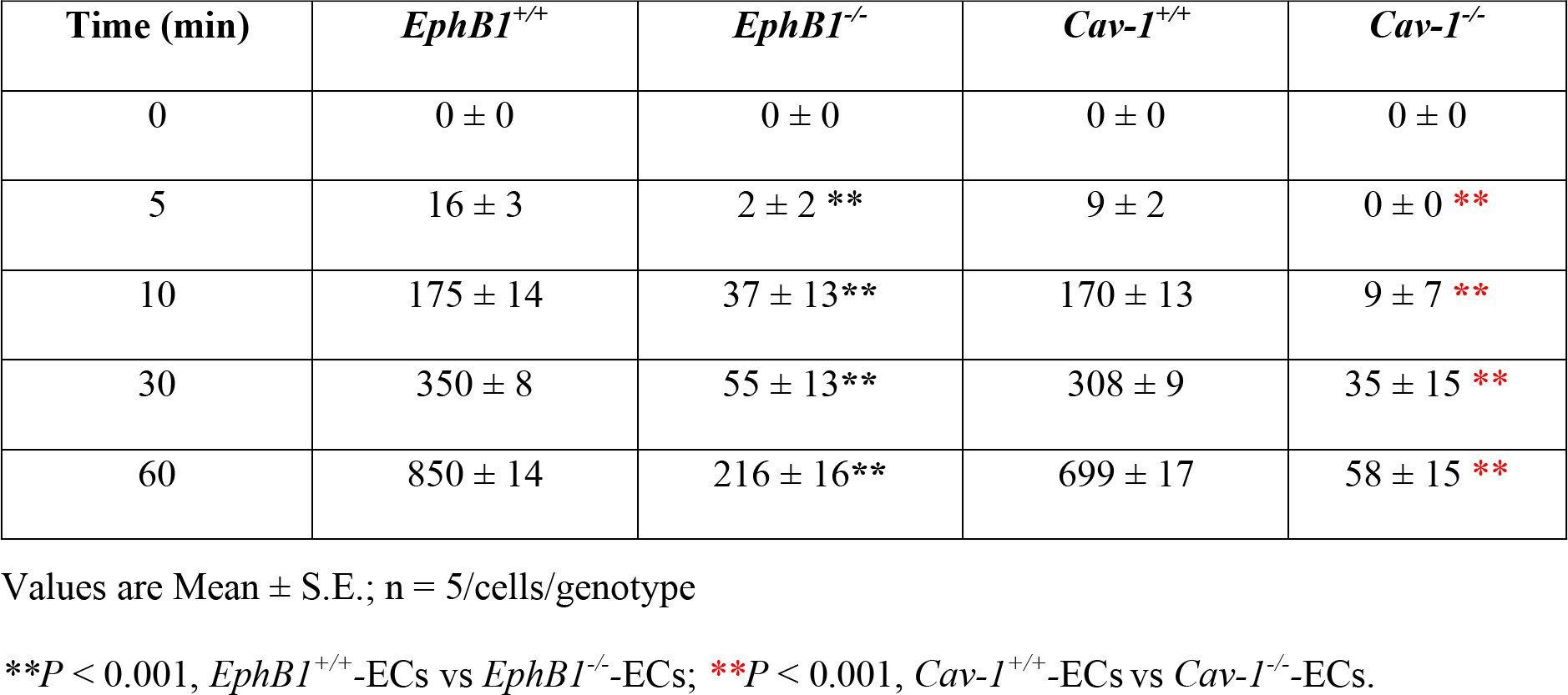
Albumin tracer particles/EC determined by 3D-SIM super resolution microscopy.

**Figure 5.**
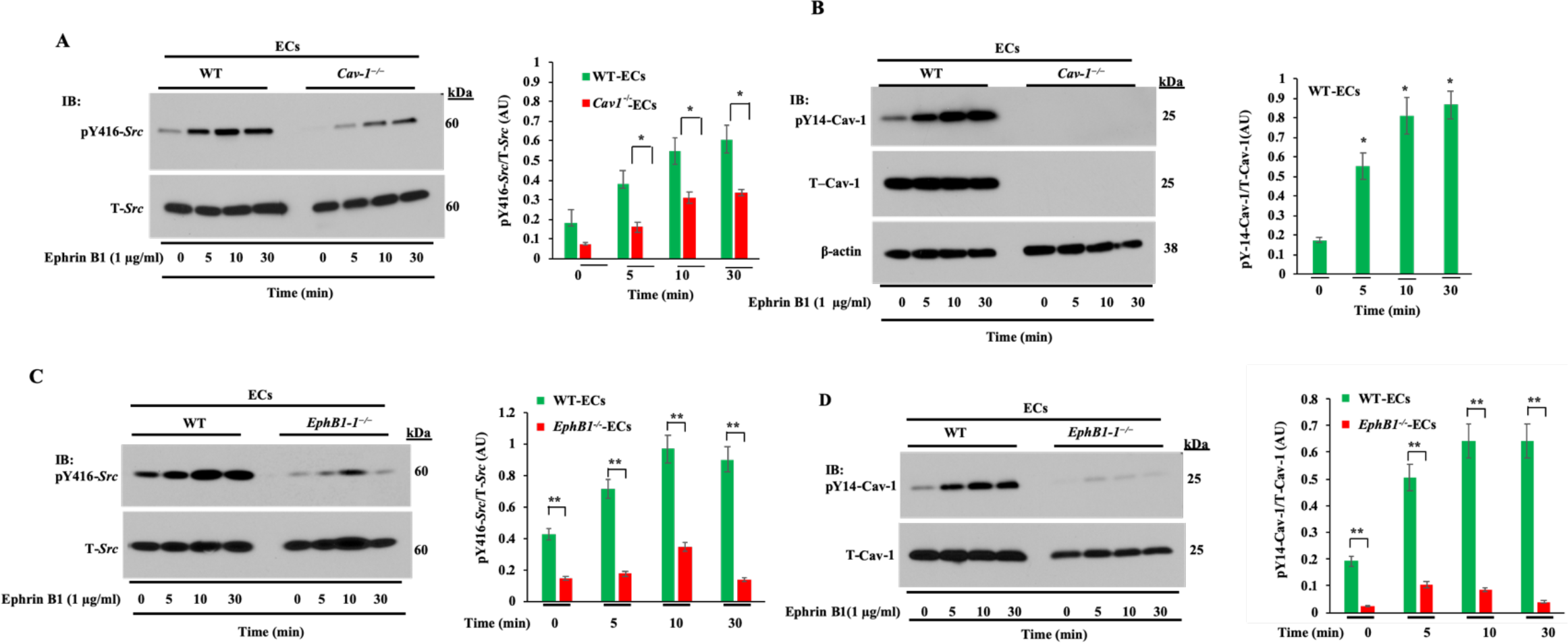
EphB1 and Cav-1 interaction is required for *Src* activation and Cav-1 phosphorylation on Y-14. **(A-D)** ECs from *Cav-1*^+/+^ (WT), *Cav-1*^−/−^, *EphB1*^+/+^ (WT), and *EphB1*^−/−^ mice exposed to EphrinB1 (EphrinB1-Fc; 1 µg/ml) for the indicated times were used for immunoblotting to determine phosphorylation of *Src* on Y-416 (*left panels*) and Cav-1 on Y-14 (pY14-Cav-1) (*right panels*). Blots shown are representative of 3 to 4 separate experiments. Values are shown as mean ± S.E. *p< 0.05; **p< 0.001; Compared with controls; WT-LECs vs *Cav-1*^−/−^ or *EphB1*^−/−^.

**Figure 6.**
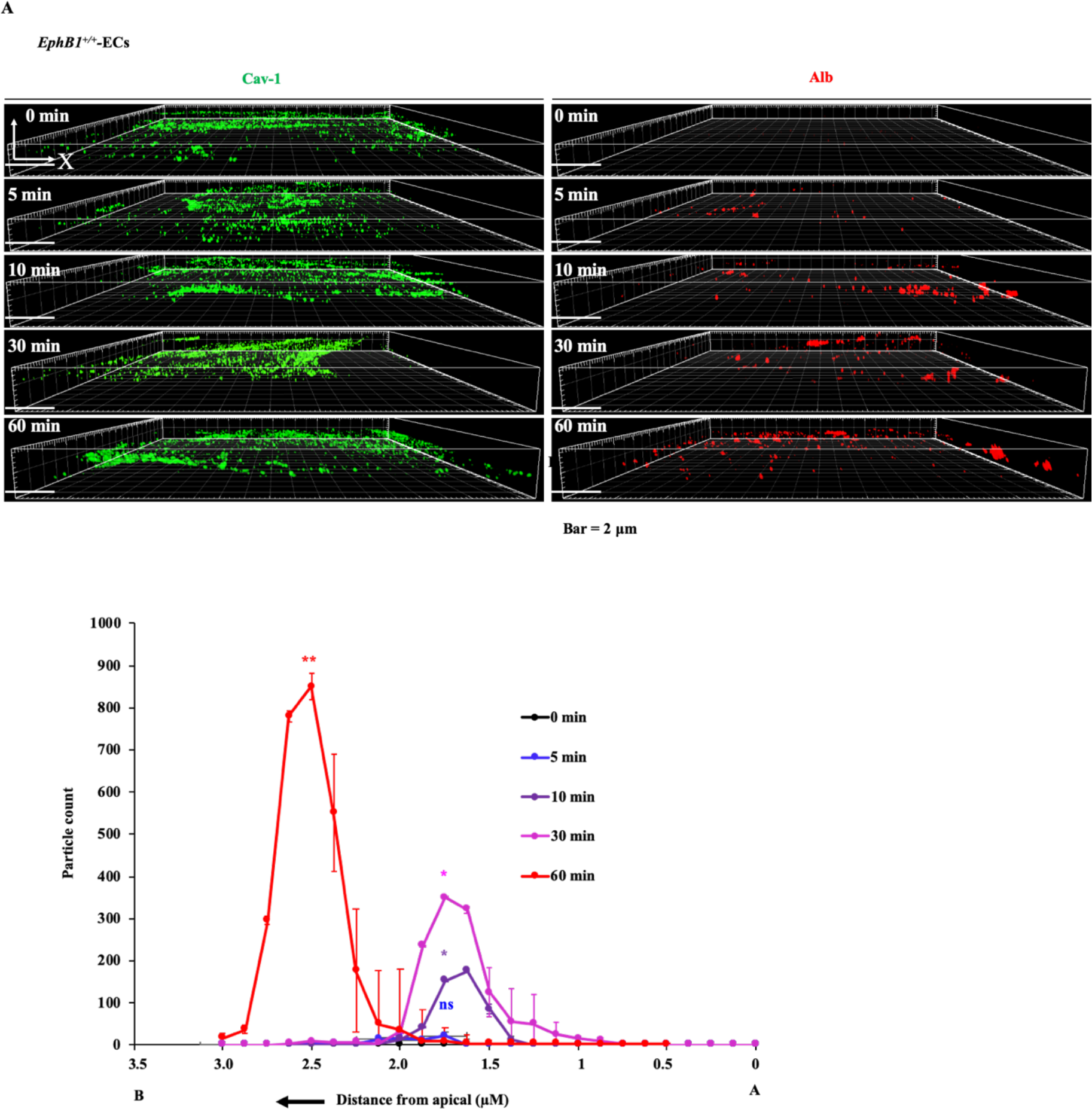
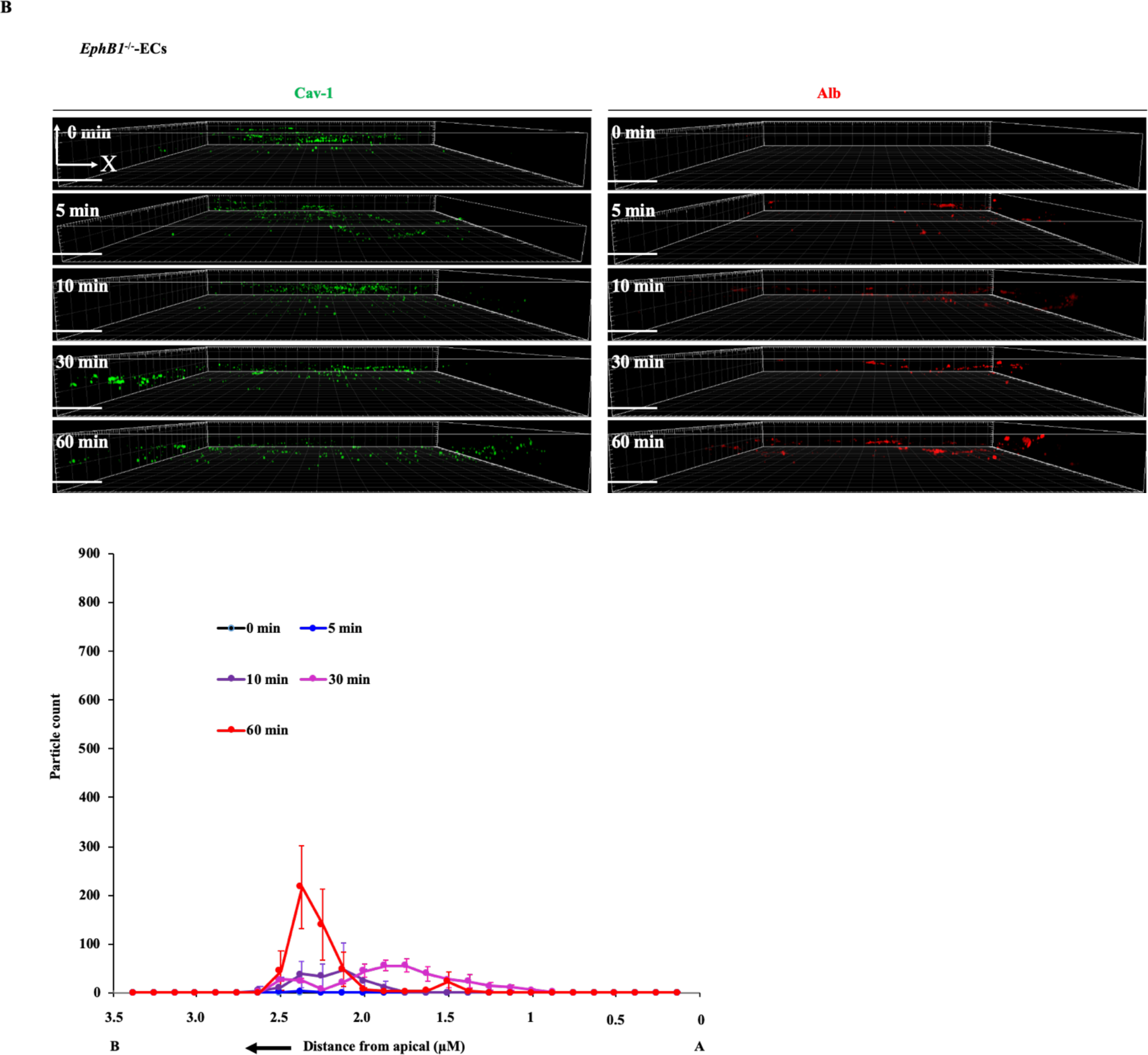
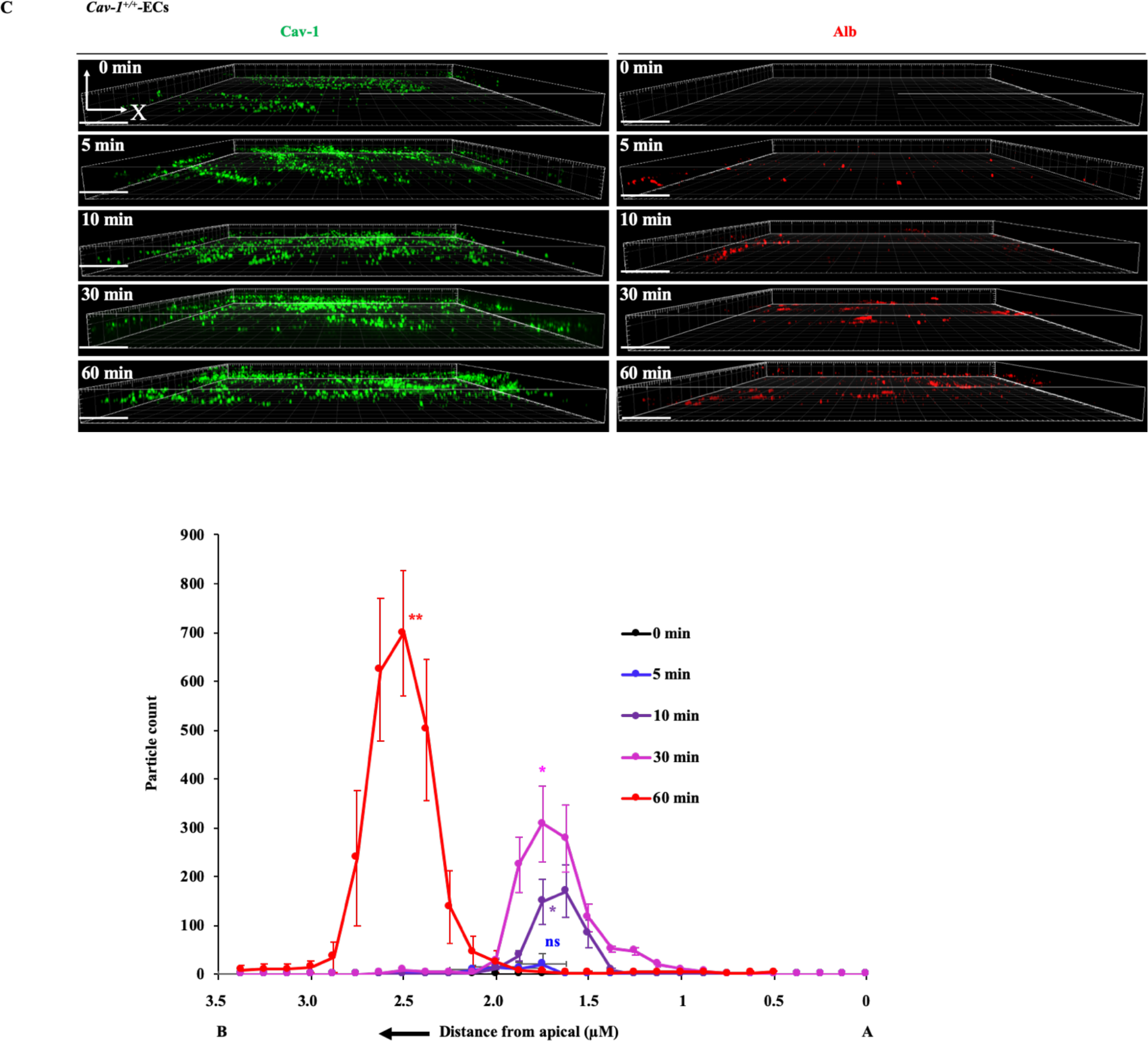
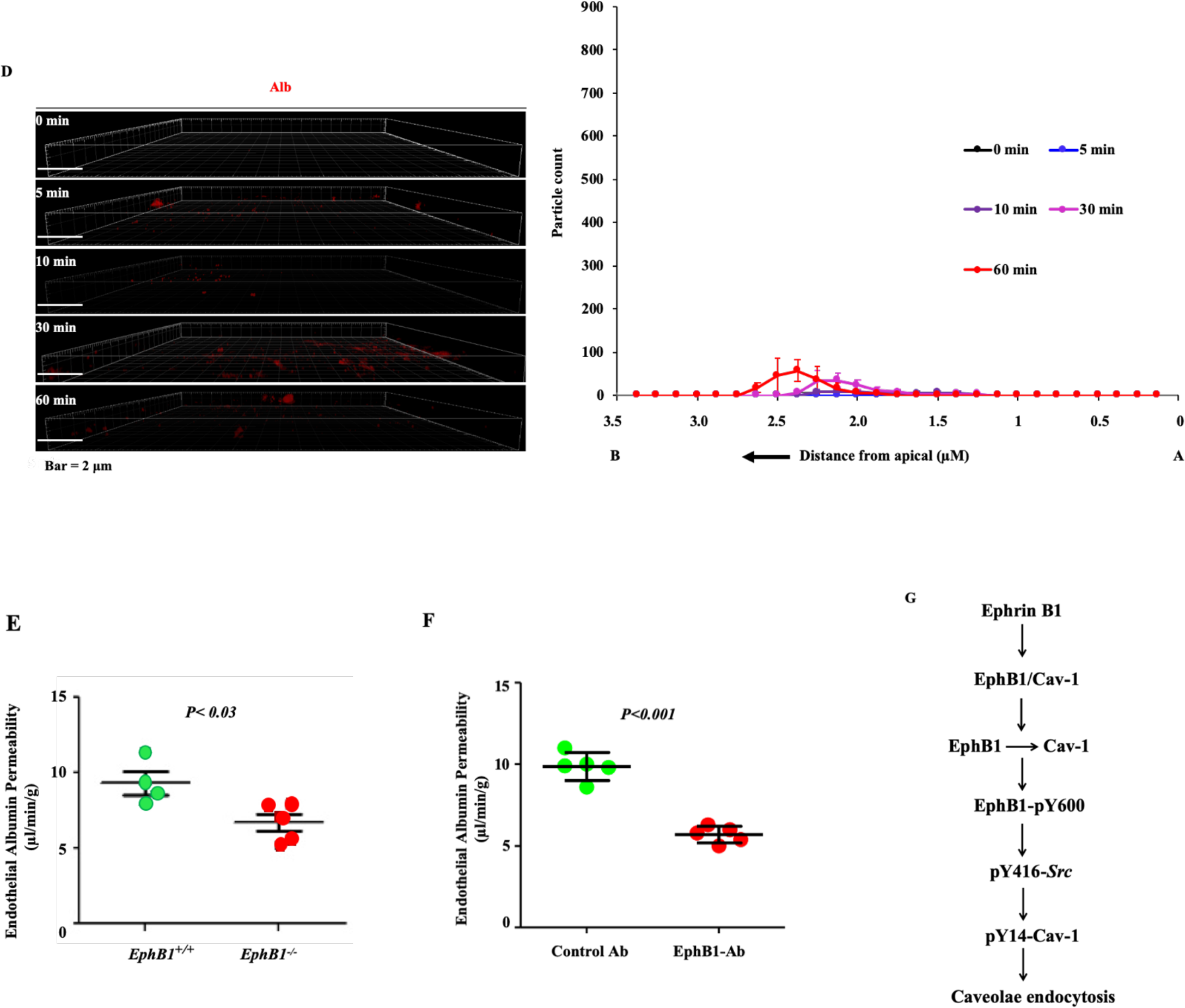
(A-D) EphB1 regulates caveolae-mediated endocytosis. **(A)** WT-ECs (EphB1^+/+^); **(B)** *EphB1^−/−^*-ECs; (**C**) Cav-1^+/+^-ECs; (**D**) *Cav-1*^−/−^-ECs. (**A-D**) Representative 3D images of single cell labeled with Cav-1 (green) and albumin tracer (red) tracking from apical and basal aspects of ECs are shown. Image in each 3D-SIM data set was processed using the Particle Analysis function in Image J software. Results show time course of Ephrin B1-Fc (1 μg/ml) -induced albumin internalization and passage from apical to basal aspects of ECs (**A-C**, *bottom panels* and **D** *left panel*). A, apical side; B, basolateral side. *p< 0.05; **p< 0.001; Compared with 0 min. (**E and F) EphB1 deletion impairs endothelial flux *in vivo*.** Flux of tracer ^125^I-albumin was measured in wild type (*EphB1*^+/+^) and *EphB1*^−/−^ mice. N = 5 per group **(E)**. Anti-EphB1 pAb (5µg/ml) similarly reduced albumin transcytosis compared to control antibody (**F**). N = 5 per group (*EphB1*^+/+^). (**G**) Model for EphB1/Cav-1 signaling-mediated caveolae endocytosis in ECs.

We also addressed the *in vivo* relevance of EphB1/Cav-1 interaction in mediating endocytosis in endothelia using tracer ^125^I-albumin (44). *EphB1*^−/−^ mice showed a marked reduction in endothelial ^125^I-albumin uptake as compared to WT (Fig. 6E). Treatment with an anti-EphB1 pAb receptor blocking antibody significantly reduced ^125^I-albumin uptake (Fig. 6F), indicating that EphB1 is required for caveolar endocytosis *in vivo*.

## Discussion

In the present study, we have uncovered a functionally important association of the receptor tyrosine kinase EphB1 with Cav-1 in endothelial cells in coordinately regulating the activation of *Src* and *Src* mediated phosphorylation of Cav-1 on Y14 in response to activation of EphB1. We showed that CSDBM of EphB1 is constitutively bound to the scaffold domain of Cav-1 (CSD) masking the EphB1 kinase domain, and thus holding *Src* activity in abeyance. However, the activation of EphB1 by its native ligand Ephrin B1 uncoupled EphB1 from Cav-1, exposing the EphB1 kinase domain to activate *Src* and *Src*-dependent phosphorylation of Y-14 Cav-1, a phospho-active site on Cav-1 (21–23) that signaled endocytosis of caveolae.

Multiple signaling proteins have been shown to bind CSD like EphB1 described in the present study (24–33). However, it is argued on the basis of structural and bioinformatic analysis that physical interactions between CSD and CSDBM may be less common than believed since CSDBM is buried in many of the CSDBM containing proteins and thus inaccessible to bind CSD (45). CSD sequence is embedded in the lipid bilayer (45, 46) that may also interfere CSD/CSDBM-dependent interactions (45,46). However, studies in which caveolae were reconstituted to determine binding of CSD and CSDBM containing proteins showed clear evidence of binding of some proteins, *Src* kinases (*Src, Fyn*) and dynamin-2 to non-phosphorylated Cav-1 and TRAF2 to Y-14 phosphorylated Cav-1 (47). Our results based on multiple lines of evidence including using heterologous expression system and imaging showed that EphB1 binds the CSD of the non-phosphorylated Cav-1 via the EphB1 CSDBM.

The canonical Ephrin ligand/Eph receptor interaction mediates conformational changes in Eph receptor which can activate *Src* tyrosine kinase (1,2). Phosphorylation of the conserved Y-600 in the cytoplasmic juxta-membrane region of EphB1 is essential for the recruitment and activation of *Src* (36). Our 3D-SIM super-resolution imaging and FRET studies showed that the ligand Ephrin B1 induced dissociation of EphB1 from Cav-1. We observed that the uncoupling unmasked the EphB1 kinase domain (which also contains CSDBM) to auto-phosphorylate EphB1 at Y-600 and induce *Src* kinase activation and subsequently Y-14 Cav-1 phosphorylation. Co-expression of mutant EphB1^Y600F^ with WT-Cav-1 failed to induce phosphorylation of *Src* on Y416 and Cav-1 on Y-14. These findings thus show that EphB1/Cav-1 dissociation-mediated *Src* activation is a key switch responsible for phosphorylation of Y-14 Cav-1, the signal responsible for activating caveolae-mediated endocytosis (21–23). While the basis of *Src* phosphorylation is not clear, a tenable mechanism may involve a conformation change induced by Ephrin B activation of EphB1 (1,2) that forces the uncoupling of EphB1 from Cav-1 and activation of *Src* kinase.

Cav-1 is essential for caveolae biogenesis in endothelial cells as evident most prominently by studies in which deletion of Cav-1 prevented the formation of caveolae (16,19,20). CSD was key for caveolae biogenesis since caveolae did not form when CSD is deleted (46). In the present study, we observed that deletion of CSDBM on EphB1 also disrupted the formation of caveolae. That both Cav-1 and EphB1 induced the same caveolae biogenesis defect suggests that Ephrin B1/Cav-1 interaction through CSD/CSDBM is a critical requirement for the formation of caveolae. An important question is why other proteins interacting with CSD of Cav-1 such as eNOS and *Src* do not influence the genesis of caveolae as was the case with Ephrin B1/Cav-1 interaction. Unlike EphB1, their loss did not disrupt caveolae biogenesis (48). We showed that the transmembrane EphB1 is basally associated with Cav-1 at the plasma membrane unlike eNOS and *Src* (29,32). The membrane association of Cav-1 and EphB1 may thus stabilize Cav-1 and facilitate the formation of homo-oligomers or hetero-oligomers with Cav-2 at the plasmalemma, which are important determinants for caveolae generation (13,14,49).

We demonstrated that *Src-*dependent Cav-1 phosphorylation on Y-14 is essential for endocytosis via caveolae in ECs (21–23). To address how EphB1 induced activation of *Src* in caveolae microdomains leads to phosphorylation of Cav-1, we determined Ephrin B1-induced *Src* activation and phosphorylation of Y-14 Cav-1 in ECs of *EphB1*^−/−^ mice. We observed that Ephrin B1-induced *Src* activation and Cav-1 phosphorylation were prevented in ECs of *EphB1*^−/−^ mice. Consistent with these results, caveolae-mediated endocytosis was also abrogated in ECs of *EphB1*^−/−^ mice. These findings thus indicate that EphB1 signaling plays a pivotal role in regulating Cav-1 function responsible for caveolae-mediated endocytosis in ECs.

The uptake of albumin in the endothelium via caveolae has remained a vexing question. Reports have suggested the existence of albumin binding proteins that promote binding and internalization of plasma albumin in caveolae and caveolar trafficking by a “pump” mechanism (50,51). The high ambient plasma albumin concentration of 3-4g% suggests that such a mechanism would be fully saturated and constitutively active, precluding any homeostatic control of interstitial oncotic pressure that is governed by the plasma albumin concentration; thus, the role of caveolae in trafficking macromolecular cargo such as albumin in the endothelium seems untenable (12, 45, 47). We showed that the ligand of EphB1, Ephrin B1, itself induced *Src* phosphorylation on Y-14 Cav-1 and thereby activated the caveolar uptake and transport machinery. Our results thus suggest that EphB1 ligation by Ephrin B released into the plasma (1,9,52) or Ephrin B from neighboring cell at the sites of cell-to-cell contact (5–7) may activate caveolar dynamics and transport via caveolae through the activation of *Src* and *Src*-mediated Y-14 phosphorylation on Cav-1.

## Material and Methods

### Antibodies and other reagents

Anti-Cav-1 rabbit polyclonal (cat # 610059) and anti-phos-Y14-Cav-1 mouse monoclonal (cat # 611338) antibodies were from BD Transduction (San Jose, CA). Anti-*Src* rabbit monoclonal (cat # 21235), anti-phos-Y416 *Src* rabbit polyclonal (cat # 21015), and pan anti-ubiquitin polyclonal (cat # 3933S) antibodies were obtained from Cell Signaling (Danvers, MA). Anti-cavin-1 (PTRF) pAb (cat #18892-1) and Anti-cavin-2 (SDPR) pAb (cat #12339-1) were from ProteinTech group (Chicago, IL). Chicken polyclonal β-galactosidase antibody (cat #ab9361) was from Abcam. Rabbit polyclonal von Willebrand Factor antibody (cat #AB7356) obtained from Millipore Corp. Rabbit polyclonal antibody against EphB1 sequence (AA 528-541, DDDYKSELREQLPL) (anti-EphB1 pAb) was custom produced by Sigma-Aldrich (St. Louis, MO). Anti-EphB1 mouse monoclonal against EphB1 sequence (AA 528-541, DDDYKSELREQLPL) was custom made by GenScript. N-terminal biotin labeled cell permeable antennapedia peptide (AP; RQIKIWFQNRRMKWKK); Cav-1 scaffold domain peptide (DGIWKASFTTFTVTKYWFYR) as a fusion peptide to the C terminus of the N-terminal biotin labeled AP; and EphB1 panning (receptor antagonistic) peptide (EWLSPNLAPSVRGSGSK) and scrambled control peptide (RTVAHHGGLYHTNAEVK) were synthesized by GenScript. PCR primers were custom synthesized from Integrated DNA technologies. Mouse recombinant EphrinB1-Fc chimera (cat # E0653-200UG) and bovine serum albumin (cat #05470-1G) with purity >98 % were from Sigma. Alexa Fuor-594 BSA (cat #A13101), Alexa Fuor-488 chicken anti-rabbit (cat #A21441), and Alexa Fluor-647 goat anti-mouse (cat #A21236) were from Life Technologies (Carlsbad, CA). Tracer ^125^I-human serum albumin (HSA) was from AnazaoHealth Corporation (Tampa, FL).

### Mice

EphB1-deficient (*EphB1*^−/−^) and EphB1-tc (EphB1-βgal fusion receptor lacking the tyrosine kinase and C-terminal domains) mice generated on a CD1 background (3,4). Caveolin-1 deficient (*Cav-1*^−/−^) mice on a C57BL/6J background (19) were from Jackson Labs (Farmington, CT). Age-matched *EphB1*^+/+^, *EphB1*^−/−^, *EphB1-tc*, *Cav-1*^+/+^ and *Cav-1*^−/−^ littermates were used for all experiments. All mice were housed in the University of Illinois Animal Care Pathogen Free Facility in accordance with institutional guidelines and guidelines of the US National Institute of Health. Veterinary care of these animals and related animal experiments was approved by the University of Illinois Animal Resources Center.

### Quantitative real-time PCR

Total RNA was isolated from lung tissue was reverse-transcribed with oligo(dt) primers and SuperScript reverse transcriptase (Invitrogen). The cDNA obtained was mixed with SYBR Green PCR mix (AB Applied Biosystems). An ABI prism 7000 was used for quantitative PCR. GAPDH expression served as an internal control. The following primers were used: mouse Cav-1α Forward, 5’-AATACGTAGACTCCGAGGGACA-3’, and reverse, 5’-GACCACGTCGTCGTTGAGAT-3’; mouse Cav-1β, forward, 5’-TGAACTTTTCTTCCCACCGCT-3’, reverse, 5’-TCAAAGTCAATCTTGACCACGTC-3’; GAPDH forward, 5’-ACCCAGAAGACTGTGGATGG-3’, and reverse, %’-CACATTGGGGGTAGGAACAC-3’.

### Expression constructs and Transfection

C-terminal CFP-tagged Cav-1 was prepared as described previously (23). pcDNA3 vector expressing mouse EphB1 C-terminal YFP-tagged (WT-EphB1-YFP), C-terminal YFP-tagged EphB1^Δ808-815^ (EphB1^Δ808-815^-YFP), and C-terminal YFP-tagged EphB1-^Y600F^ (EphB1-^Y600F^-YFP) were custom prepared by GenScript. DNA sequencing was performed to verify all the expression constructs sequences. COS-1 cells plated on 60 mm dishes at 60% confluency were transfected with WT-Cav-1-CFP (1 µg) plus WT-EphB1-YPF (2.5 µg), WT-Cav-1-CFP (1 µg) plus EphB1^∆808-815^-YFP (2.5 µg), or WT-Cav-1 (1 µg) plus EphB1-^Y600F^-YFP (2.5 µg using Superfect transfection reagent (cat #301305, Qiagen). Media was replaced 6 h after transfection with fresh DMEM media containing 10% FBS. After 72 h, cells were harvested and lysed in radioimmunoprecipitation assay (RIPA) buffer containing protease and phosphatase inhibitor cocktails. For 3D-SIM imaging experiments, 24 h after transfection cells plated on High Tolerance Coverslips (pcs-170-1818, MatTek) and at 72 h, cells were used for experiments.

### Live-cell fluorescence resonance energy transfer (FRET) imaging

Live-cell FRET imaging was performed as in (23).

### Endothelial cells

Human lung microvascular endothelial cells (HLMVECs) and Endothelial growth media-2 (EGM-2) were purchased from Lonza (Walkersville, MD, USA). Lung endothelial cells (ECs) from mice were isolated with mAb to the adhesion molecule CD31 (PECAM-1) (53).

### Immunoblotting

Endothelial cells were washed three times with phosphate-buffered saline (PBS) at 4°C and lysed in lysis buffer (50 mM Tris-HCl, pH7.5, 150 mM NaCl, 1 mM EGTA, 1% Triton X-100, 0.25% sodium deoxycholate, 0.1% SDS, 10 µM orthovanadate, and protease-inhibitor mixture) (54). Mouse tissues were homogenized in lysis buffer (54). EC lysates or tissue homogenates were resolved by SDS-PAGE on a 4–15% gradient separating gel under reducing conditions and transferred to a Duralose membrane. Membranes were blocked with 5% dry milk in TBST (10 mM Tris-HCl pH7.5, 150 mM NaCl, and 0.05% Tween-20) for 1 h RT and then incubated with the indicated primary antibody diluted in blocking buffer overnight at 4°C. For phospho-specific blots, membranes were incubated overnight at 4°C with the primary antibody diluted in TBST containing 5% bovine serum albumin. Next, membranes were washed three times and incubated with appropriate HRP-conjugated secondary antibody. Protein bands were detected by enhanced chemiluminescence.

### Immunostaining

Cells grown on High Tolerance Coverslips (pcs-170-1818, MatTek) were incubated with serum free medium (5 mM HEPES/HBSS, pH 7.4) for 2 h and stimulated with Ephrin B1-Fc (1 μg/ml) for indicated time periods. Cells were washed, fixed with 3% paraformaldehyde in HBSS, permeabilized with 0.1% Tween-20 in 5 mM HEPES/HBSS, pH 7.4, and blocked with 5% human serum diluted with 5 mM HEPES/HBSS, pH 7.4 (blocking buffer) for 2 h at room temperature. After blocking, cells were incubated with primary antibodies (1 μg/ ml), anti-EphB1 mouse mAb or anti-Cav-1 rabbit mAb in blocking buffer overnight at 4°C. Cells were washed and incubated with 1 μg/ml secondary antibody Alexa Fluor-647 goat anti-mouse and Alexa Fluor-488 chicken anti-rabbit for 1 h. After washing coverslips were mounted on a microscope slides with prolong gold-antifade reagent (Invitrogen).

### Immunoprecipitation

Cell lysate (150 μg protein) was subjected to immunoprecipitation. Each sample was incubated overnight with 1 μg/ml of the indicated antibody at 4°C. The next day, protein A/G beads were added to the sample and incubated for 1 h at 4°C. Immunoprecipitates were then washed three times with wash buffer (Tris-buffered saline containing 0.05% Triton X-100, 1 mM Na_3_VO_4_, 1 mM NaF, 2 μg/ml leupeptin, 2 μg/ml pepstatin A, 2 μg/ml aprotinin, and 44 μg/ml phenylmethylsulfonyl fluoride). Immunoprecipitated proteins were used for immunoblotting.

### 3D-Structured Illumination Microscopy (SIM)

All 3D-SIM imaging was performed on a DeltaVision OMX SR system (GE) equipped with an Olympus 60X/1.42 NA objective and refractive index matched immersion oil (n=1.516-1.518) at room-temperature. Full-frame structure illuminated image sequences were taken at 125 nm Z-axis sections for multiple Z positions; the same exposure and excitation parameters were undertaken to avoid pixel saturation and maintain the validity of comparison across samples. 3D-SIM images were reconstructed using Softworx (Applied Precision) offline using a Wiener filter coefficient of 0.001. Reconstructed 3D-SIM data was further processed by Imaris software (Bitplane, Zurich, Switzerland). To ensure colocalization accuracy, all color channels in the 3D-SIM system have been aligned using standard samples. The image in each 3D dataset was processed using the Particle Analysis function in ImageJ to quantify the number and intensity of specific protein particles in each Z section.

### Transmission Electron Microscopy (TEM)

WT and *EphB1*^−/−^ mice were anesthetized with ketamine/xylazine (100 mg kg/5 mg kg) by i.p. injection. Harvested organs were perfused with HBSS containing EM fixative (2% glutaraldehyde, 3% paraformdehyde, 0.1 M sodium cacodylate, pH 7.2). Tissues blocks (1 × 2 mm) prepared fixed in fresh fixative, rinsed in 0.1 M sodium cacodylate, post-fixed in 1% OsO_4_ in 0.1 M sodium cacodylate, rinsed, stained en block with Kellenberger’s uranyl acetate in water, dehydrated through graded ethanol, and embedded in LX-112 resin using propylene oxide. Ultrathin sections of 20-40 nm were cut and mounted on grids, stained with uranyl acetate and lead citrate and examined under a Joel 1220 electron microscope (55).

### Caveolae endocytosis assay

Caveolae-mediated endocytosis of tracer albumin was measured as described (21,22). Briefly, endothelial cells grown on High Tolerance Coverslips (pcs-170-1818, MatTek) were washed, and incubated for 2 h in serum-free medium, followed Alexa Fluor 594-conjugated albumin (0.1 mg/ml Alexa Fluor 594-BSA mixed into 2 mg/ml of non-fluorescent BSA) in 5 mM HEPES-buffered HBSS at 37°C for 30 min (21,22). After this period of incubation, cells were washed, 3% paraformaldehyde in HBSS, permeabilized with 0.1% Tween-20 in 5 mM HEPES/HBSS, pH 7.4, and blocked with 5% human serum diluted with 5 mM HEPES/HBSS, pH 7.4 for 2 h at RT. After blocking, cells were incubated with anti-Cav-1 rabbit mAb (1 μg/ml) in blocking buffer overnight at 4°C. Cells were washed and incubated with 1 μg/ml secondary antibody Alexa Fluor-488 chicken anti-rabbit for 1 h. After washing coverslips were mounted on a microscope slides with prolong gold-antifade reagent (Invitrogen).

### Endothelial permeability assay

We determined transendothelial permeability in lung vessels in WT or *EphB1*^−/−^ mice were anesthetized using 2.5% sevoflurane in room air, and 2 µCi of ^125^I-labeled albumin tracer injected intravenously according to an approved animal protocol. At 30 min after tracer injection, 100 µl blood sample withdrawn from a vein to determine blood tracer counts. Organs were then cleared of circulating tracer by whole-body perfusion via the right heart using RPMI supplemented with 3% unlabeled albumin and lungs were excised and counted for γ-radioactivity. Transendothelial albumin permeability was calculated in ml/min/g dry tissue from blood and tissues counts, and values were normalized to tissue dry weight as described (44). To study the effects of the anti-EphB1 pAb on transendothelial albumin permeability, we used the method as described by us (53, 56). Murine lungs were perfused via the pulmonary artery with RPMI solution containing 3% unlabeled albumin (2 ml/min, 37°C). All preparations underwent a 20-min equilibration perfusion, and then received control Ab or anti-EphB1 pAb via a side-arm of the pulmonary artery cannula to achieve a final perfusate concentration of 5 µg/ml each. The ^125^I-albumin tracer was infused via a separate side-arm for a 30 min-period. A perfusate sample was collected to determine blood tracer counts and the washed out for 6 min, a period sufficient to reduce effluent counts to background levels. Lungs were counted for gamma radioactivity, and transendothelial albumin permeability was quantified in units of ml/min/g dry lung.

### Statistical analysis

Results were analyzed by an unpaired two-tailed Student’s *t*-test. Differences in mean values were considered significant at a p value <0.05.

## ACKNOWLEDGEMENTS

This work was supported by National Institutes of Health Grants R01 HL-128359, R01 GM-117028, R01 HL-122157, and P01 HL-060678.

## AUTHORS CONTRIBUTIONS

C.T. and A.B.M. designed the research. C.T., S.C.R., D.M.W., and S.M.V. performed most experiments. S.C.R., P.T.T., and G.C.H.M involved in 3D-SIM imaging. R.V.S. and R.D.M. involved in EM experiments. M.H. generated EphB1 knockout and EphB1-tc mice. C.T. analyzed and interpreted the results. C.T. and A.B.M. wrote the manuscript.

