## Supplementary Information for "EphB1 in Endothelial Cells Regulates Caveolae Formation and Endocytosis"

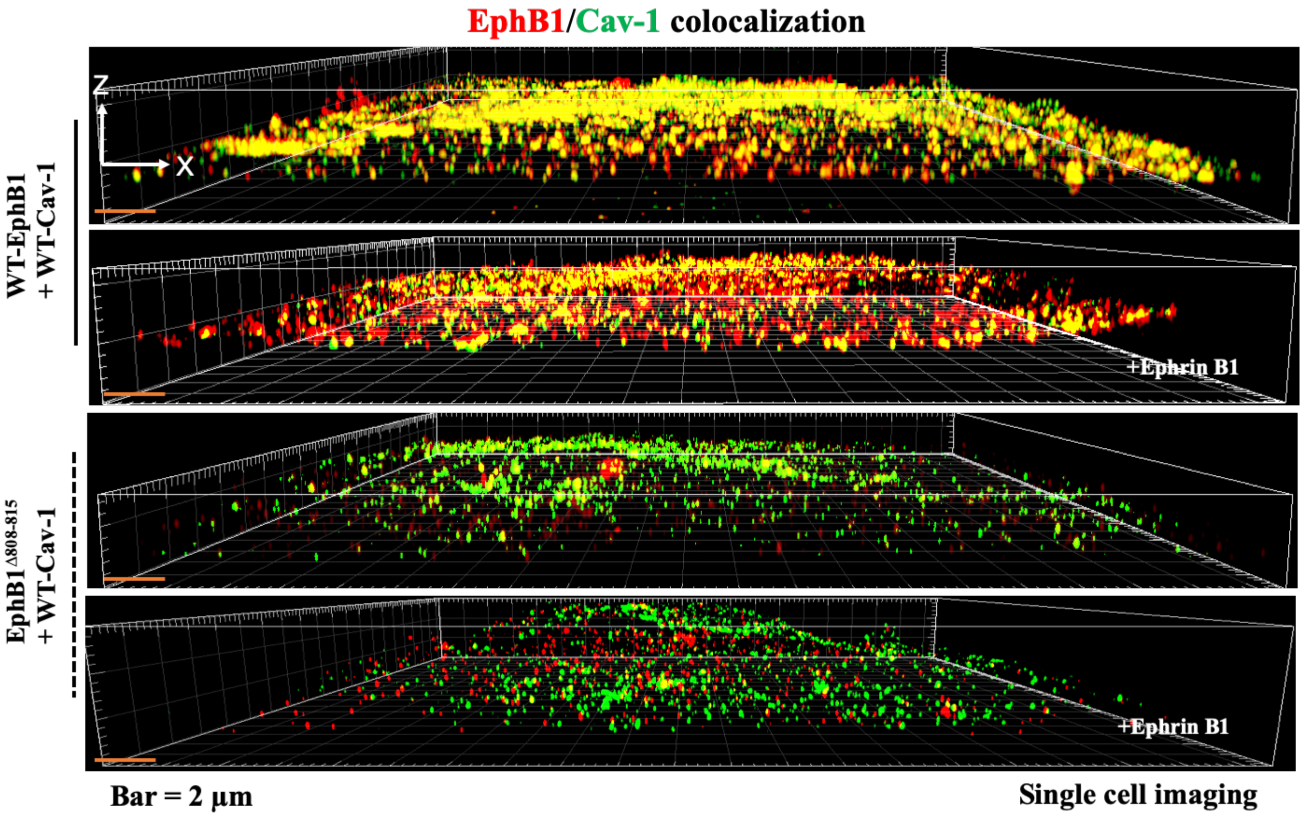


**Supplemental Figure 1A.** COS-1 cells transfected with WT-EphB1 + WT-Cav-1 or EphB1$^{\Delta 808\text{-}815}$ + WT-Cav-1 were used for 3D-SIM imaging. Representative 3D image of single cell from 3D-SIM showing EphB1 interacts with WT-Cav-1 (*top panel*). Ephrin B1-Fc (1 µg/ml; 10 min) challenge caused dissociation of EphB1 from Cav-1 (*second panel*). *Third and fourth panels*, representative 3D image of single cell from 3D-SIM showing CSDBM deleted EphB1 (EphB1$^{\Delta 808\text{-}815}$) failed to interact with WT-Cav-1. Ephrin B1-Fc stimulation had no significant effect on this interaction.

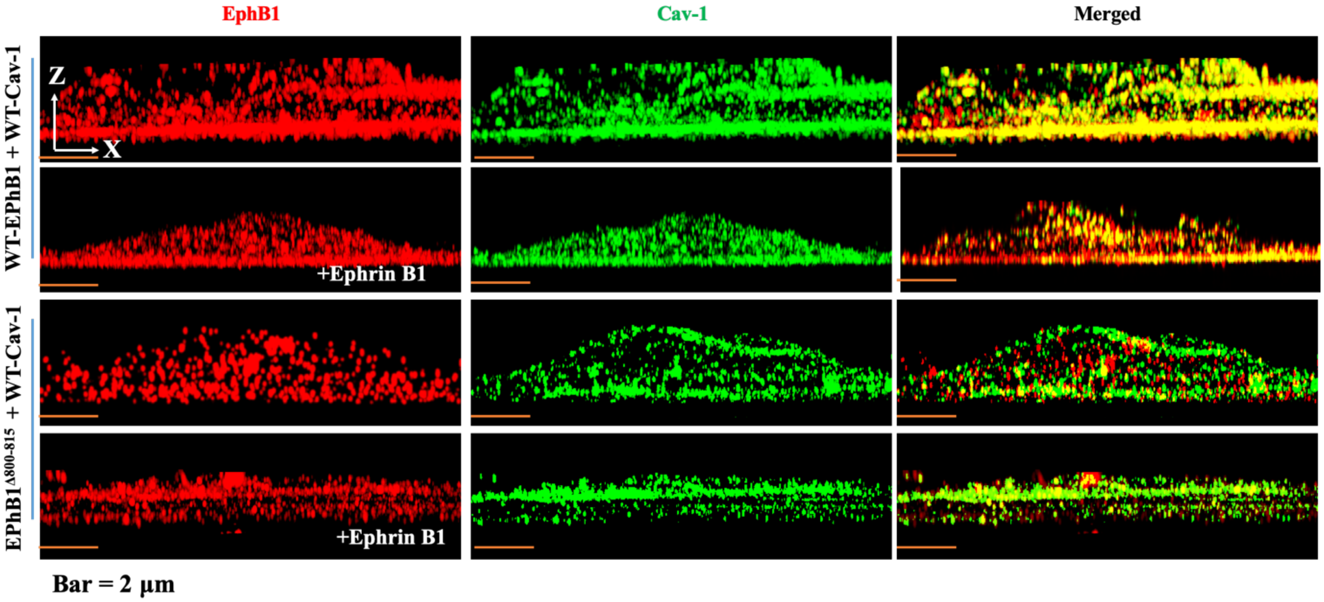


**Supplemental Figure 1B.** COS-1 cells were transfected with WT-EphB1 + WT-Cav-1 or EphB1$^{\Delta808\text{-}815}$ + WT-Cav-1 used for 3D-SIM imaging. Representative sagittal plane view of single cell 3D-SIM image showing EphB1 interacts with WT-Cav-1 (*top panel*). Ephrin B1-Fc (1 µg/ml; 10 min) challenge caused dissociation of EphB1 from Cav-1 (*second panel*). *Third and fourth panels*, representative sagittal plane view of single cell 3D-SIM showing CSDBM deleted EphB1 (EphB1$^{\Delta808\text{-}815}$) failed to interact with WT-Cav-1. Ephrin B1-Fc stimulation had no significant effect on this interaction.

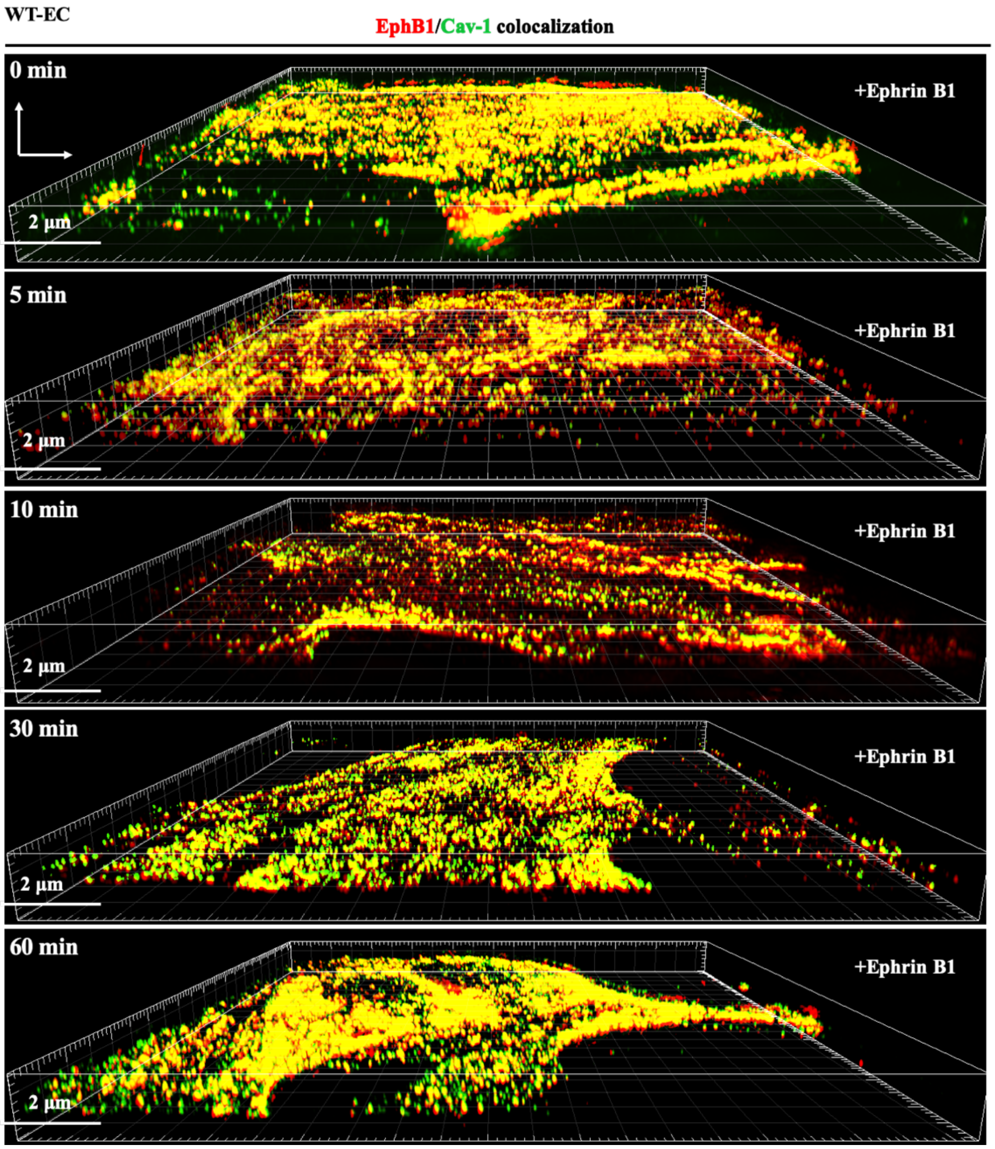


**Supplemental Figure 1C.** 3D image of single WT-EC from 3D-SIM showing of colocalization of EphB1 with Cav-1. Ephrin B1-Fc stimulation caused the dissociation of EphB1 and Cav-1 and then EphB1 re-associated with Cav-1 in EC.

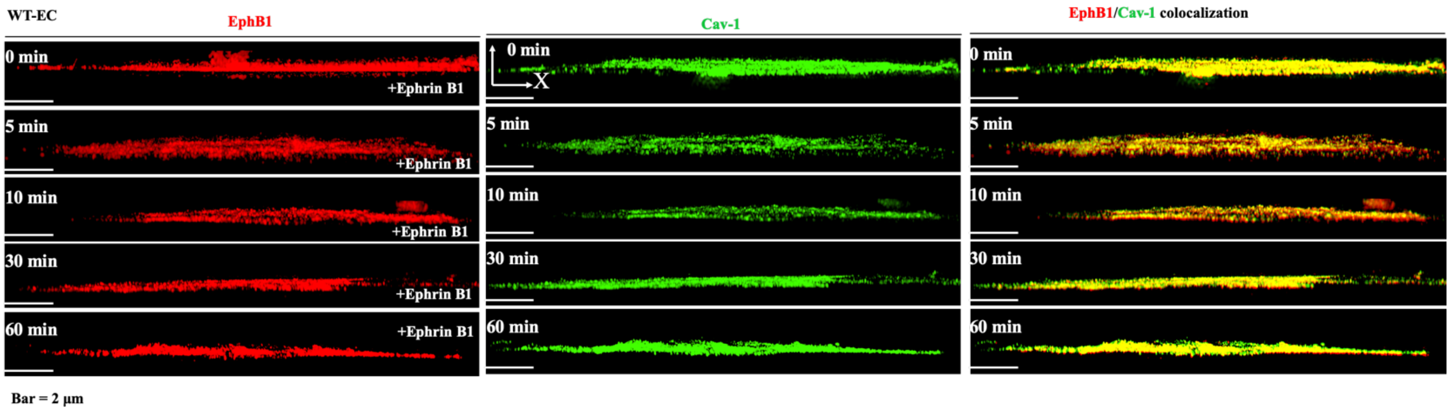


**Supplemental Figure 1D.** Sagittal plane view image of single WT-EC from 3D-SIM image showing colocalization of EphB1 with Cav-1. Ephrin B1-Fc stimulation caused the dissociation of EphB1 and Cav-1 and then EphB1 re-associated with Cav-1 in EC.

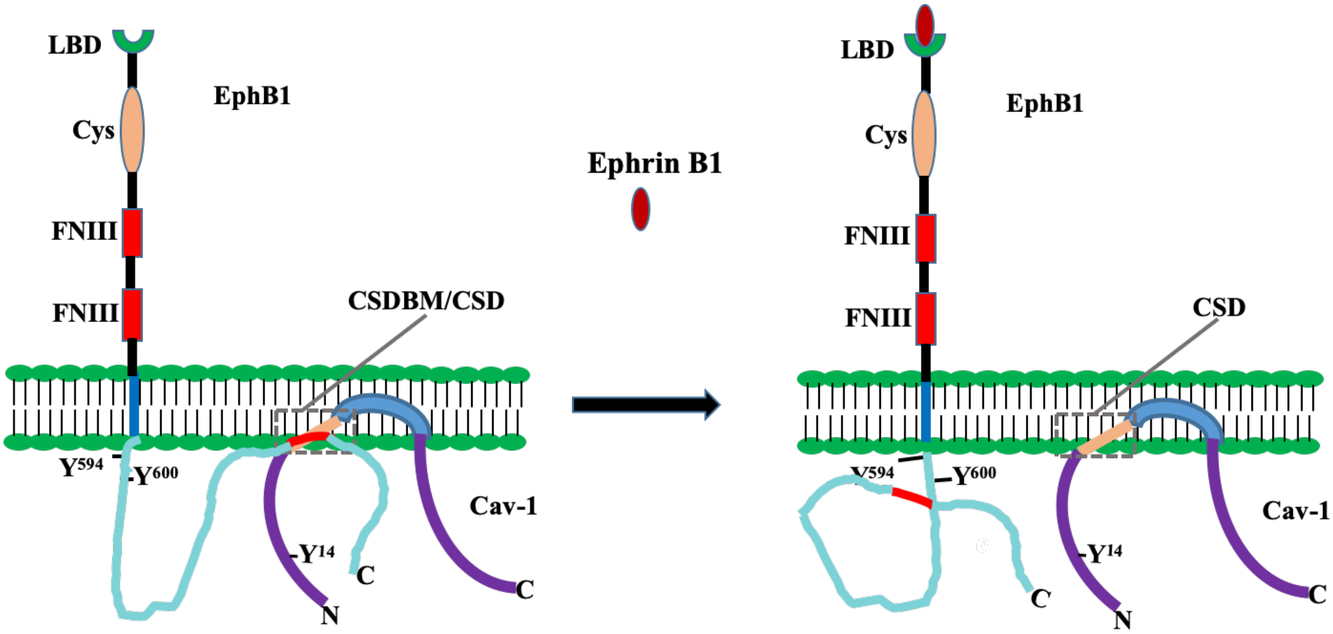


**Supplemental Figure 1E.** Model for the interaction between EphB1 and Cav-1 in ECs based on our results.

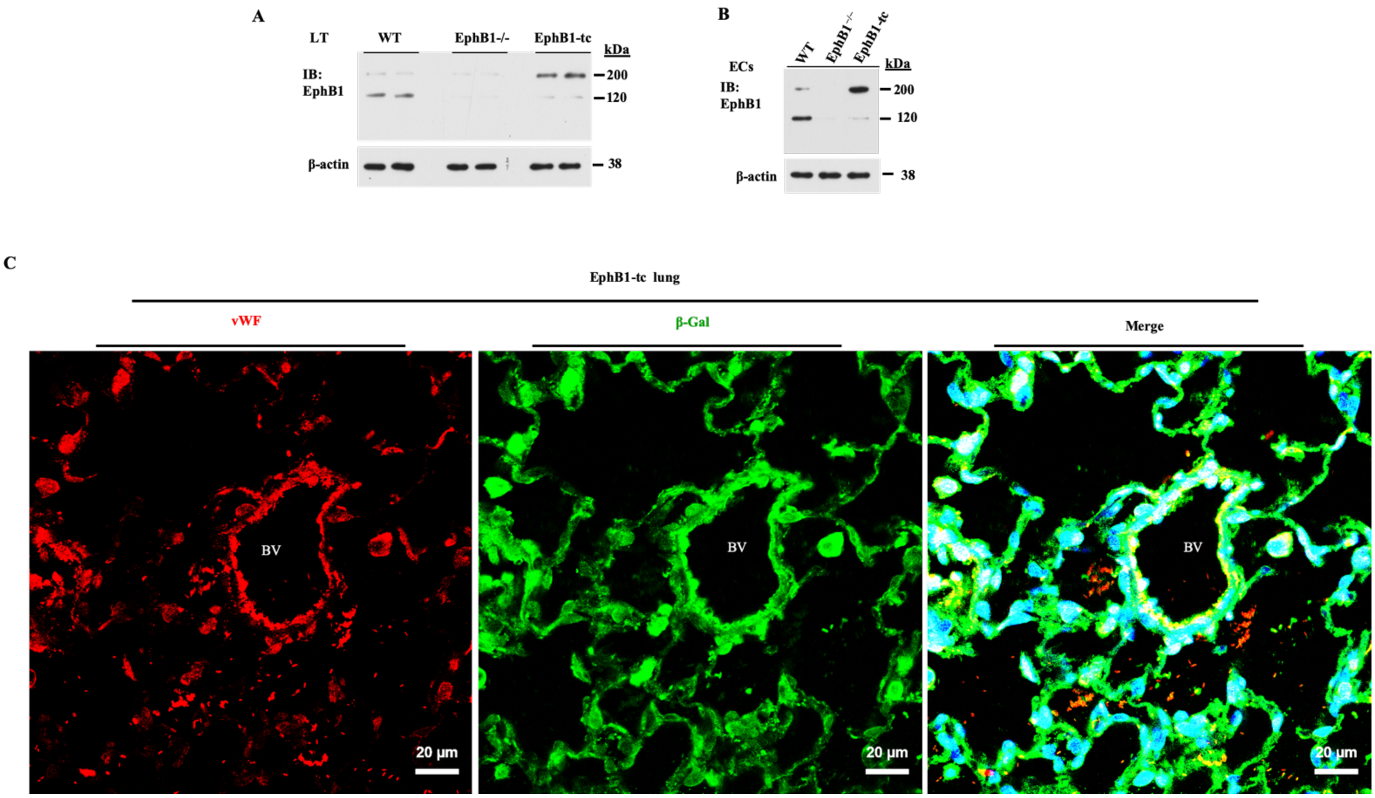


**Supplemental Figure 2. Expression of EphB1 in adult lungs. (A, B)** Lungs and lung ECs from WT (CD1), *EphB1*$^{-/-}$, and *EphB1-tc* (EphB1-βgal fusion) mice were used for Western analysis to determine EphB1 expression. (C) Immunostaining of lung sections from EphB1-tc mice with EC marker vWF and β-galactosidase (β-Gal) to assess EphB1 expression lung vascular ECs.
